# Deeplasmid: Deep learning accurately separates plasmids from bacterial chromosomes

**DOI:** 10.1101/2021.03.11.434936

**Authors:** William B Andreopoulos, Alexander M Geller, Miriam Lucke, Jan Balewski, Alicia Clum, Natalia Ivanova, Asaf Levy

**Affiliations:** Joint Genome Institute, US Department of Energy, LBNL Berkeley, California, USA; National Energy Research Scientific Computing Center (NERSC), Berkeley, California, USA; Department of Plant Pathology and Microbiology, The Institute of Environmental Science, The Robert H. Smith Faculty of Agriculture, Food, and Environment, The Hebrew University of Jerusalem, Rehovot, Israel; Department of Computer Science, San Jose State University, CA, USA

## Abstract

Plasmids are mobile genetic elements that play a key role in microbial ecology and evolution by mediating horizontal transfer of important genes, such as antimicrobial resistance genes. Many microbial genomes have been sequenced by short read sequencers and have resulted in a mix of contigs that derive from plasmids or chromosomes. New tools that accurately identify plasmids are needed to elucidate new plasmid-borne genes of high biological importance. We have developed Deeplasmid, a deep learning tool for distinguishing plasmids from bacterial chromosomes based on the DNA sequence and its encoded biological data. It requires as input only assembled sequences generated by any sequencing platform and assembly algorithm and its runtime scales linearly with the number of assembled sequences. Deeplasmid achieves an AUC-ROC of over 93%, and it was much more precise than the state-of-the-art methods. Finally, as a proof of concept, we used Deeplasmid to predict new plasmids in the fish pathogen *Yersinia ruckeri* ATCC 29473 that has no annotated plasmids. Deeplasmid predicted with high reliability that a long assembled contig is part of a plasmid. Using long read sequencing we indeed validated the existence of a 102 Kbp long plasmid, demonstrating Deeplasmid’s ability to detect novel plasmids.

**Availability:** The software is available with a BSD license: <u>deeplasmid.sourceforge.io</u>. A Docker container is available on DockerHub under: billandreo/deeplasmid.

## Introduction

Plasmids are ubiquitous extrachromosomal elements capable of semi-autonomous replication and transmission between microbial host cells. Typically, bacterial plasmids are small (<80 kb) circular replicons. Natural plasmids often carry a cargo of “accessory genes” that confer beneficial traits to the microbial host, such as antibacterial resistance (1, 2), bacteriophage defense (3, 4), heavy metal tolerance (5), virulence (6, 7), or unique catabolic pathways (8), thereby improving bacterial adaptation to dynamic environments. Some plasmids carry toxins and antibiotic resistance genes and thereby constitute a serious threat to human health (9). Finally, plasmids are involved in plant-microbe interactions; for instance, the nodulation plasmids of rhizobia guide the symbiosis of bacteria with plants (10). Plasmid transmission by conjugation provides an efficient mechanism of horizontal gene transfer and facilitates the spread of accessory genes in bacterial populations and communities. Therefore, the studies of plasmid genetics, evolution, and dynamics in bacterial populations have many wide-reaching practical applications, such as clinical management of antibiotic resistance (2), development of the industrial strains of bacteria for bioremediation (3) and biofertilization (4). In addition, identification of new plasmids may guide the discovery of novel antibiotic resistance genes, toxins, and genes directly involved in host-microbe interactions, and may be used as new tools for efficient gene cloning and exogenous protein expression.

Advances in genomic sequencing technologies have enabled high-throughput sequencing of genomes of microbial isolates and environmental populations (through metagenome sequencing), including their respective plasmidomes - the total collection of encoded plasmids (5). Identification and classification of plasmid sequences in this treasure trove of genomic and metagenomic data can provide a unique opportunity to study the mechanisms of plasmid persistence, transmission, and host specificity, as well as the flow of accessory genes. However, *in silico* identification of plasmid contigs in whole-genome shotgun sequences (WGS) is challenging. The challenge derives from poor genome assembly that leads to numerous plasmid-size contigs that are difficult to characterize as derived from plasmid or chromosomes. In addition, there is a limited number of high-quality, completely sequenced reference plasmids that can be compared to while annotating new genomes (6). Sequences from plasmids occasionally integrate into chromosomes, making it difficult to computationally characterize contigs from these chromosomes as plasmids or chromosomes. Further, sometimes plasmid genes have features resembling those of essential chromosomal genes (11).

A variety of *in silico* methods assisting with separation of plasmid sequences from chromosomal contigs have been developed. Some of them target a subset of plasmids mostly of clinical relevance, such as PlasmidFinder/pMLST (12) for detection and typing of plasmids from *Enterobacteriaceae* and selected Gram-positive strains. Other tools, such as PLACNET (13) rely on a combination of reference genomes and manual curation to restructure an assembly graph and separate putative plasmid contigs from those of chromosomal origin. plasmidSPAdes (14), cBAR (15), PlasFlow (16), Recycler (17), and PlasmidSeeker (18) are fully automated and perform identification of putative plasmid contigs in genome assemblies by analyzing the topology and read coverage of an assembly graph (Recycler and plasmidSPAdes) or DNA composition of assembled contigs (cBar and PlasFlow). Recycler works on paired-end reads and detects circular plasmids by leveraging assembly graphs from conventional assembly tools to assemble circular sequences likely to be plasmids (17). HyAsP starts from raw reads and combines read depth with GC content, as well as reference-based occurrences of known plasmid genes in the assembly (19). An assessment of methods that assemble plasmids from short reads (20) concluded that their accuracy is reliant on a difference in the coverage of plasmids and chromosomes; for some assemblies they demonstrated close to a 90% precision of plasmid finding with just 55% recall, whereas for assemblies with 80% recall generally the false positive rate increases by 20% (20). Moreover, most of these tools were not tested for their ability to detect novel plasmids that are experimentally validated following the computational prediction.

Existing tools have limitations due to their reliance on the circularity of the topology, bias towards certain taxonomies used in training (e.g. in PlasmidFinder) and coverage of a de Bruijn assembly graph constructed from k-mers found in reads (e. g. Recycler and plasmidSPAdes). The two software packages: cBAR (15) and PlasFlow (16), satisfy the above criteria, since they utilize only two types of data: assembled sequences themselves (PlasFlow) and features extracted from assembled sequences (cBar). PlasFlow relies on a deep neural network to find hidden structures encoded in the assembled sequences, while cBar finds plasmids by applying self-organizing maps (SOMs) to the extracted features in the form of pentamer profiles of contigs and scaffolds. The two methods also differ in the way their models are trained: PlasFlow was developed as a tool for finding plasmids in metagenome data and is pre-trained on sequence fragments of up to 10 kb long, since metagenome assemblies are typically very fragmented. In contrast, cBAR’s model is based on the pentanucleotide profiles of full-length sequences of known plasmids and chromosomes. While both cBAR and especially PlasFlow demonstrated superior performance in comparison to other methods of plasmid identification (16).

Our goal was to develop a tool for post-assembly identification of complete plasmids and plasmid-derived contigs, which (i) has high accuracy, (ii) is not biased towards the sequences of certain topology or taxonomic origin, and (iii) is able to run on genome assemblies from either short-read or long-read sequencing technologies without assembly graph or coverage information. We also confirmed that inclusion of genes and other functional features in addition to DNA sequence composition is helpful for contig and scaffold classification. We present a new Deep Learning (DL)-based method, Deeplasmid, for identification of plasmid contigs and scaffolds in WGS assemblies of microbial isolate genomes, which achieves an AUC-ROC of 93% on a sixfold cross-validation. Our method relies on a combination of assembled sequences and extracted features, including GC content (21), oligonucleotide composition, hits to plasmid- or chromosome-specific genes, as well as gene density within the contig. Since it does not require raw read data, assembly graph or coverage information, it can be applied to assembled WGS data, including shotgun metagenomic data, generated by any sequencing platform and assembly algorithm. We describe our Deeplasmid model, the training and testing methodology, and show that it is capable of automated detection of plasmid sequences with over 94% accuracy. Deeplasmid surpasses the accuracy of other tools largely due to its use of discriminating gene and protein features. We compare the accuracy of our trained model on large plasmid-containing microbial test datasets against the alternative tools cBAR and PlasFlow, thus achieving a better precision and comparable recall. Finally, we used a whole genome sequencing project of a specific microbe, *Yersinia ruckeri* ATCC 29473, applied Deeplasmid and predicted a novel plasmid in this strain. We then performed a sequencing experiment to validate that the new plasmid indeed exists as a separate replication unit. This led to discovery of a new plasmid in this pathogenic strain.

## Materials and Methods

### ACLAME-RefSeq training dataset

We prepared the labeled dataset based on two sources. As negative instances we used 50 different genera from the RefSeq.microbial dataset (22), from which plasmid and any mitochondrial or chloroplast sequences were removed based on their fasta header names. The training included Archaeal chromosomal sequences from 40 genera, which are found in RefSeq.archaea. As positive instances we used the ACLAME dataset (23), which contains 1,056 fully-sequenced plasmids that were manually curated by experts. ACLAME has higher-quality curation than refseq.plasmids since some of the NCBI records tagged as plasmids are mislabeled as chromosomal sequences and many entries do not represent complete records or contain sequence fragments of unknown origin (24, 25). From ACLAME we discarded 39 sequences (3.69%) that were shorter than 1 kbp or longer than 330 kbp because the scaffolds and contigs longer than 330 kb are almost invariably chromosomes or megaplasmids or chromids (genetic elements with plasmid-type replication systems, but carrying some indispensable genes (11)). We did not deal with the last two classes as they are special cases. To balance the dataset size we randomly selected 40,000 sequences from the RefSeq.microbial dataset, which is two orders of magnitude larger than ACLAME.plasmid dataset of size 1,017. The imbalance 1/40 in the sequence count in our ACLAME-RefSeq training dataset was compensated during the training by oversampling the ACLAME dataset. Data were shuffled before training.

### Input format

A single training data element consists of the label and two input words: x_seq_ - a 300bp contiguous subsequence sampled randomly from the full original scaffold sequence and x_f_ - a vector containing 16 features extracted from the full sequence, as described in Table 1. In order to ensure feature values like gene count, gene coding percentage, or sequence length are meaningful, the features are computed on the entire scaffold, and the values are copied into the x_f_ feature vectors for all 300bp sequences subsampled from the scaffold. The number (*m*) of 300bp subsequences sampled from each scaffold is proportional to the square root of the scaffold length. The number of samples per scaffold was chosen according to m=10+sqrt(seq_len/20) to ensure a fair representation of smaller and larger scaffolds, such that longer scaffolds do not overwhelm the training step. Each sample is a different x_seq_ associated with the feature vector x_f_ from the originating sequence. x_seq_ is one-hot encoded in 4 nucleotide bases. Namely, it is transformed into a binary array of size 300×4. The ‘N’-base (unknown) is encoded as four zeros. The values of x_f_ were normalized to be bound within [-1,1].

**Table 1.** Definition of 16 features per sequence

| Name | Definition | Type |
| --- | --- | --- |
| gc_content | GC content of contig | Float [0-1] |
| A(C/G/T)_longest_homopolymer | Length of longest homopolymer | Integer |
| A(C/G/T)_total_homopolymer | Total number of homopolymers of length >5 | Integer |
| hit_chromosome_proteins | Hit to chromosome proteins | Boolean 0/1 |
| hit_plasmid_proteins | Hit to plasmid proteins | Boolean 0/1 |
| hit_plasmid_ORIs | Hit to plasmid ORI | Boolean 0/1 |
| gene_count | Number of genes in scaffold | Integer |
| gene_percent | Coding percent of scaffold | Float [0-1] |
| polypeptide_aa_avg_len | Average length of amino acid sequence | Integer |
| len_sequence | Scaffold seq length | Integer |

### Input feature selection

We initially explored the predictive power of several extracted features in conjunction with existing Machine Learning tools. The particular choice of x_f_ variables shown in Table 1 was based on an initial sensitivity analysis with the Gradient Boosting Classifier, a classic ML method that produces a prediction model in the form of a mixture of decision trees. Moreover, we confirmed the relative importance of a feature by training our tool and running predictions with null values for the feature (namely, excluding the feature) and checking the impact on the error rate (as discussed in the Supp. Info).

We additionally included in x_f_three plasmid-specific and chromosome-specific features. These features are boolean (0 or 1) and indicate whether any hit is found to these sets of plasmid or chromosome-specific sequences:

1. Plasmid-specific DNA motifs: these are the origins of replication of known plasmids (26).
2. Plasmid-specific proteins: these are taken from 2,826 known plasmids listed on 2019 in the European Nucleotide Archive: https://www.ebi.ac.uk/genomes/plasmid.html, and removed any plasmids not isolated from Proteobacteria, Firmicutes, Bacteroidetes, or Actinobacteria using NCBI batch entrez function. We only kept plasmids from these four phyla as these are the most commonly sequenced and studied bacterial phyla and as a result most contigs that will be classified by our tool belong to these phyla. Some plasmidic proteins were extracted from publications (27–36) (Table S1). The final list included 136,638, 24,607, 1,163, and 15,449 plasmidic genes from Proteobacteria, Firmicutes, Bacteroidetes, and Actinobacteria respectively.
3. Chromosome-specific proteins: these are based on COGs of genes that are usually carried on chromosomes. The 61 COGs used for making this list are based on chromosomal housekeeping genes that are unclonable in high copy plasmids (37, 38). They appear in Table S1.

To reduce sequence redundancy the chromosomal and plasmid proteins were clustered by 90% identity using cd-hit with otherwise default parameters giving a representative sequence from each cluster (39).

### Output format

A neural network is a function

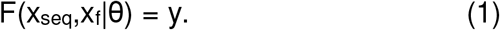

that accepts sequences of nucleotides xseq and the feature vector x_f_. The function F also depends implicitly on the DL model parameters θ, which are determined during the training process. The output of the network, Deeplasmid score y, is computed using the softmax function, which ensures that y satisfies y □ [0, 1]. By convention, the higher the score is for the sequence, the more likely it is to be a true plasmid.

### Model training

The model was trained with a binary cross-entropy loss function (40) and Adam optimizer (41). We performed supervised learning on the balanced set of 6×10^5^ data points with a batch size of 200. The initial learning rate was set to 0.001. Typically we sample 50-100 300bp long sequences per scaffold.

We used the k-fold cross-validation method, setting k=6, with five data segments merged as the ‘training’ set and one validation segment that provided the loss (model error) as feedback during training. The ‘test’ data set was hidden during the training. We performed 6 independent trainings, cycling the segments to allow each of six segments to influence a different model θ_k_. Figure 1 illustrates the k-fold training method. Each model was trained for 30 epochs, until it converged, as shown in Figure 2a.

**Fig. 1.**
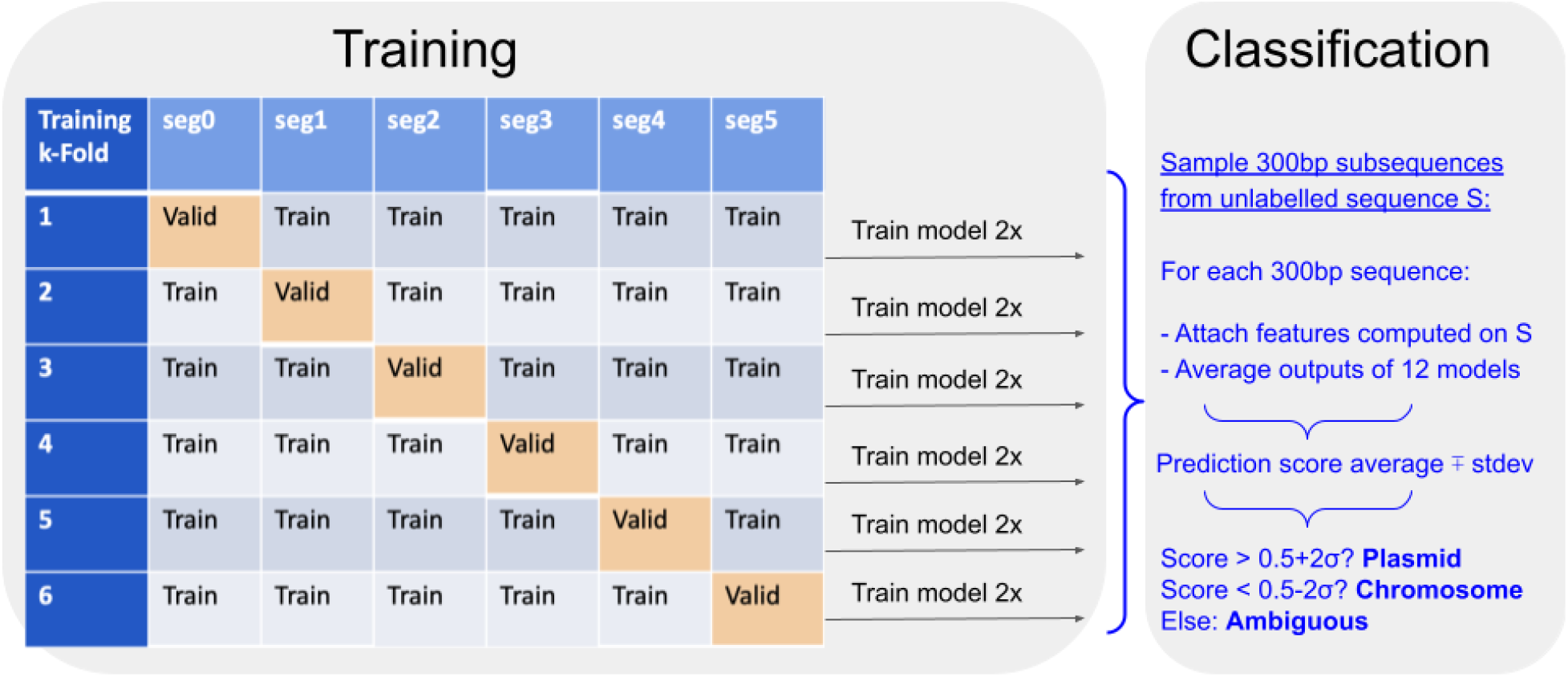
Deeplasmid training, validation and testing. The plasmid and chromosome dataset was split into six segments, of which five were used in training a model. The 6^th^segment was used for validation of the trained model. We repeated over the training twice to derive 12 different models. Using 12 models allows reducing the effects of random variance in the predictions.

**Fig. 2.**
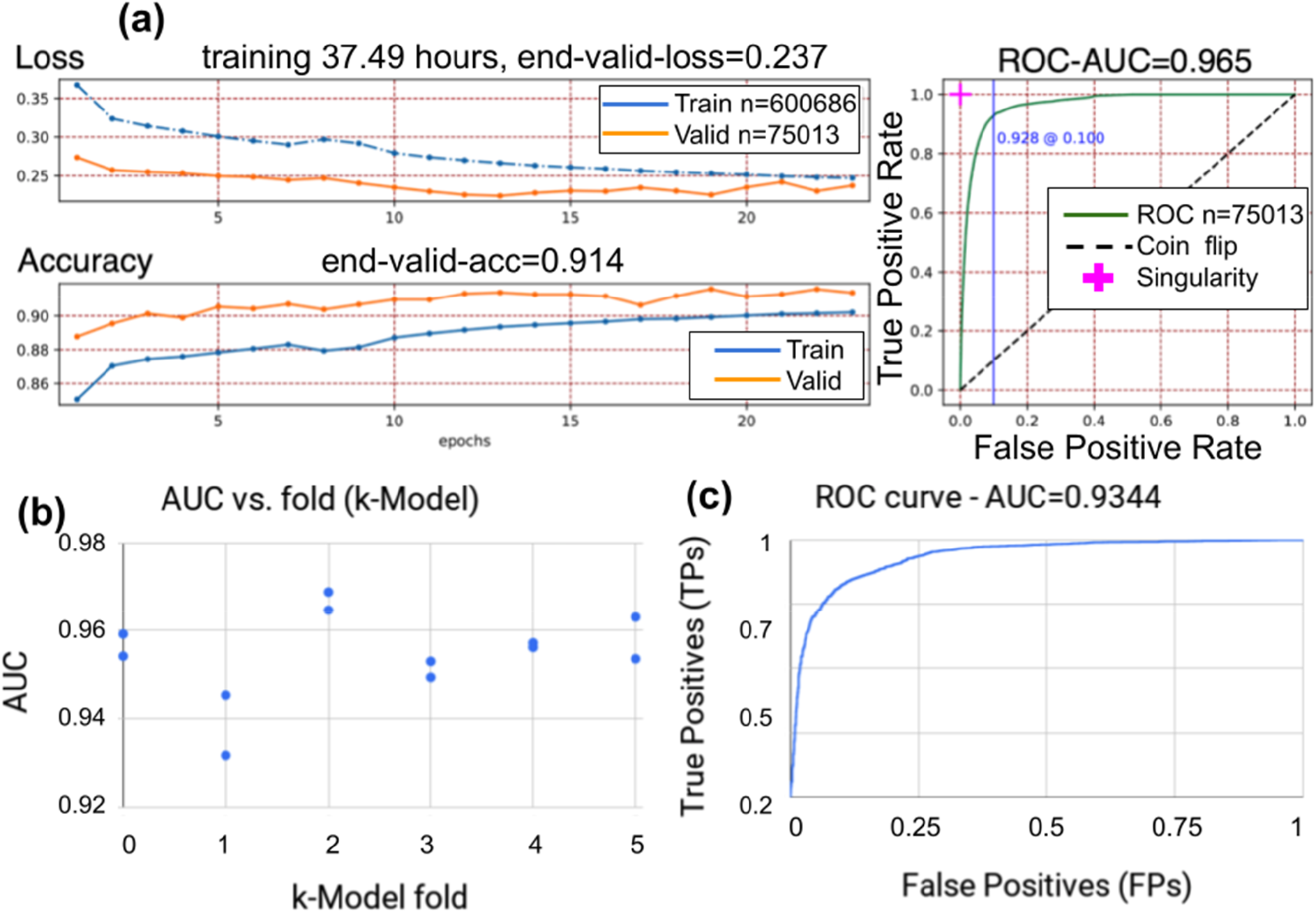
**(a)** Training convergence through the epochs. Loss and accuracy are shown as a function of epochs. The training and validation data are balanced by oversampling the plasmid class. **(b)** The training was repeated 12 times on the plasmid-chromosome dataset to derive twelve models (two per validation segment). All models achieved an accuracy (AUC) on the validation segment of over 0.93 with a small statistical variance in the prediction accuracy. **(c)** The ROC-AUC on the IMG test dataset after training the model is 0.9344.

**Fig. 3.**
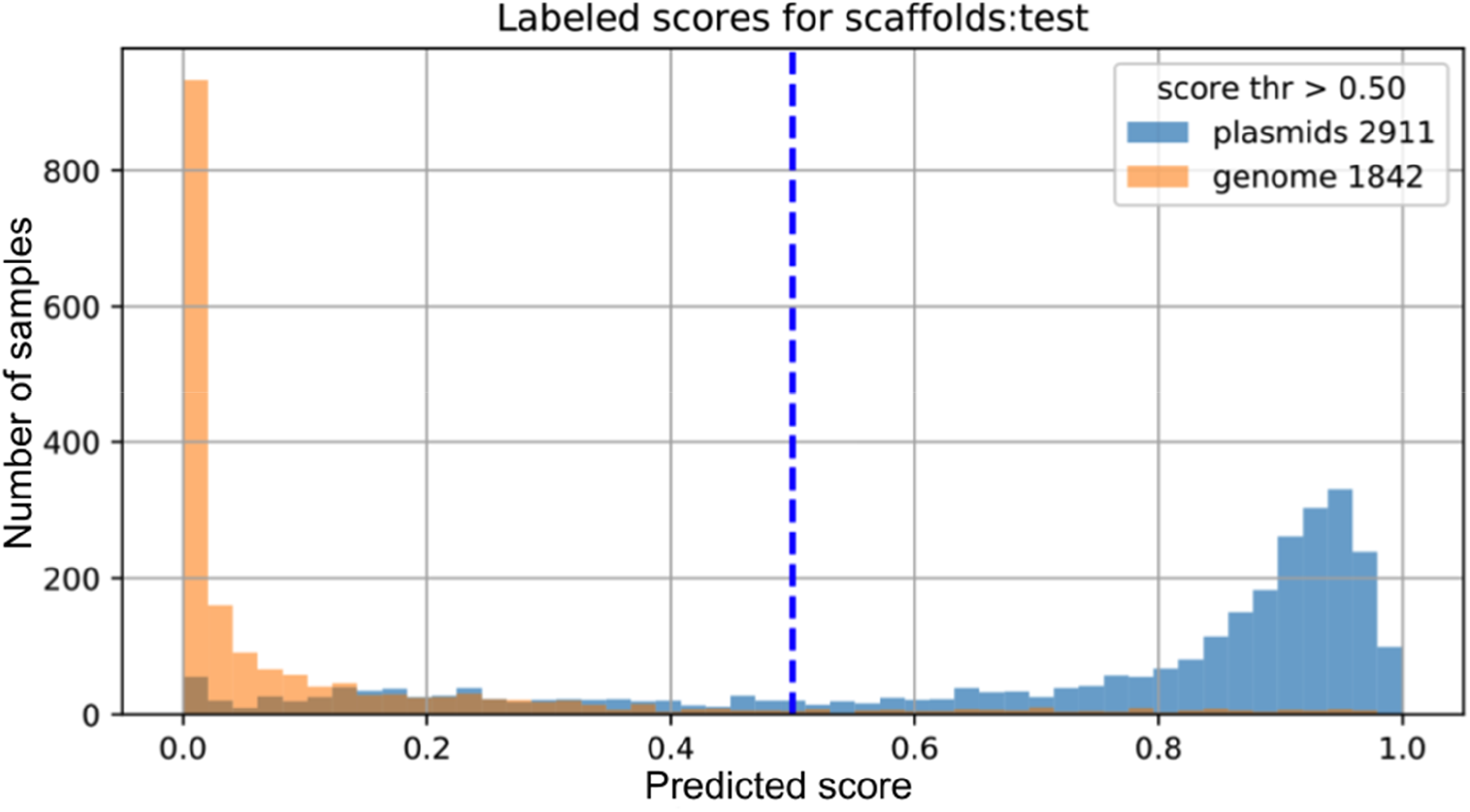
Evaluation on the IMG test dataset. The class separation is clear based on a threshold of 0.5. The percent of plasmids classified above the threshold is 2264/(2264+599+48)=77.77% (recall). The percent of chromosomes classified below the threshold is 1702/(1702+140)=92.4%.

### Model topology

We have used the LSTM-based network (42) (Figure S4) to transform the one-hot encoded nucleotide sequence into a one-dimensional vector. The left branch is made out of two LSTMs, accepts a 300bp nucleotide sequence x_seq_, and compresses information into a vector of 40 features. The right branch is fully connected, accepts the feature vector x_f_, and produces a vector of 20 features. Both outputs are concatenated and passed to another block of fully connected layers whose output is one value - the Deeplasmid score y (Eq. 1). This model was implemented in Keras (43) with Tensorflow 1.3.0 (44) as the backend. The deep learning model architecture is shown in Figure S3.

### Prediction for one 300bp sequence

For each k-fold we saved two models, resulting in 12 different saved models. This was done to reduce the effects of random variance in the predictions, as well as to ensure that the results were reproducible for each k-fold. To make a prediction on a 300bp sequence we ran the sequence through all 12 models and then average the score:

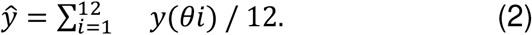

The results are shown in Table S2.

### Prediction for one scaffold

One scaffold is sampled 50-100 times and for each 300bp sequence the average score is computed as in (Eq. 2). Next, the scaffold-average score (yavr) and its standard deviation (σ) are computed. We allow for 3-way classification as “plasmid”, “chromosome”, or “ambiguous”:

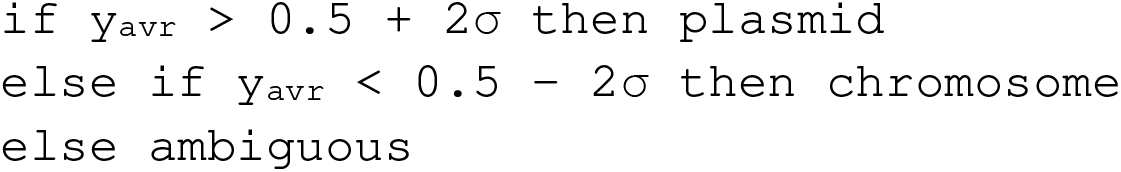

### DNA extraction and Oxford Nanopore sequencing

We validated our plasmid computational prediction using genome re-sequencing with Oxford Nanopore long read sequencing. We used strain *Yersinia ruckeri* ATCC 29473 (IMG genome ID: 2609460118), which was received as a gift from Dr. Yasuo Yoshikuni. Bacteria grew in the final volume of 2L Luria Broth until OD 600 0.3 was reached. DNA was extracted with Qiagen Genomic Tip 100/G (Cat No./ID: 10243) and Genomic DNA Buffer Set (Qiagen, Cat No./ID: 19060), by following suggested instructions. DNA concentration and quality were tested with Nanopore, Qubit and TapeStation. Prior sequencing samples were prepared with Ligation Sequencing Kit (SQK-LSK109) and Native Barcoding Expansion 1-12 (EXP-NBD104) and finally sequenced by Oxford Nanopore MinION. Reads were assembled using Canu 2.0 (45). Reads were also assembled in parallel by Shasta 0.6.0 (46). The 3,754,417 bp circular DNA and the 102,560 circular DNA were found from the Canu and Shasta assemblies, respectively. To map linear scaffolds of *Yersinia ruckeri* ATCC 29473 to the newly assembled plasmids, blastn was used, with a bitscore cutoff of 30,000. The blastn hits were visualized using DNAFeaturesViewer (47). The IMG scaffold names were shortened in Figure 4; all scaffolds displayed are prefixed by “Ga0059170_”, e.g. scaffold 114 is named “Ga0059170_114” in the IMG database. Scaffold “Ga0059170_103” coordinates 1-145,000 were mapped to the newly found plasmid (fragment is at 8 o’clock in Figure 4, marked 103*), away from the rest of the scaffold (12 to 5 o’clock in Figure 4). This subsequence is small enough to be processed by Deeplasmid and was thereby re-ran through Deeplasmid to be predicted as a separate piece of DNA. Annotation (Figure 4C) was performed based on IMG scaffold annotations (scaffolds Ga0059170_112 and Ga0059170_113).

**Figure. 4.**
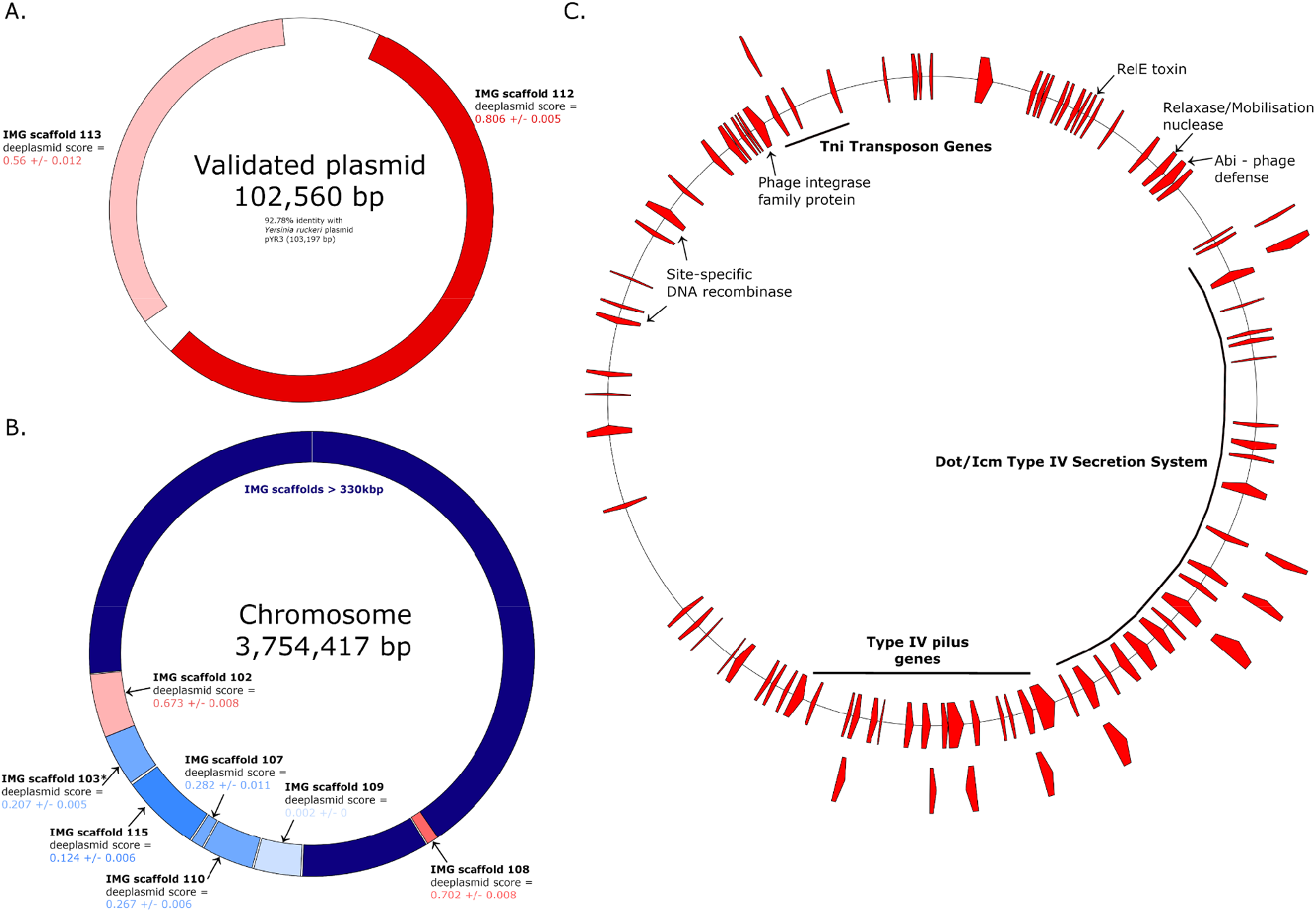
Deeplasmid vaidation. *Yersinia ruckeri* ATCC 29473 was sequenced with Oxford Nanopore MinION and assembled with Canu and with Shasta. (A) The assembled contigs included a circular piece of DNA that shares 92.78% identity with a known *Yersinia* plasmid (pYR3; Genbank: LN681230.1). Two linear scaffolds of this genome were predicted by Deeplasmid to be from plasmids (shades of red), and indeed they align with the newly-found plasmid. (B) The assembled contigs also contained a chromosome. Most of the linear scaffolds for this genome did not undergo Deeplasmid prediction, due to their large size (>330kb; dark navy blue). However, those within the size range were largely predicted to be chromosomal in origin (shades of blue), along with two misclassifications (shades of red). Scaffold 103* is a subsequence of a larger IMG scaffold; this short region was predicted by Deeplasmid here (see Materials and Methods). (C) Functions of many ORFs identified on the validated plasmid are classically associated with plasmids.

## Results

### Training and feature selection

We assembled a training dataset of bacterial and archaeal chromosomes from 90 genera that was retrieved from Refseq, and we retrieved 1,017 plasmids from ACLAME, (Materials and Methods). Features included sequence-related physical features and features related to the genetic content of the contig (Table 1, Table S1, Materials and Methods). The first feature group includes gc_content, repeats: number and size of the longest homopolymers, and total length of the sequence. The second feature group includes boolean variables of whether the sampled sequence shares similarity with chromosomal genes, plasmidic genes, or origin of replication sequences. For example, ribosomal proteins are usually carried on chromosomes and plasmid replication and mobilization genes are carried on plasmids. We also included in this group the number of genes carried on the scaffold, the percent of the scaffold which is coding, and the size of the genes (“polypeptide_aa_avg_len”) as through manual inspection we noted that plasmids tend to have more intergenic regions and smaller genes than chromosomes.

### Deeplasmid predictions for the ‘test’ data

The training dataset was divided into six segments (Materials and Methods). One segment, called ‘validation’, was used for validating a model’s training. Twelve models were trained in total, two for each selection of a validation segment. The model training over 30 epochs is characterized by an increase in prediction accuracy (defined as the ratio of correct classifications to all queries) and decrease in error (loss) on the validation segment. The Receiver Operating Characteristic (ROC) curve is shown in Fig. 2a. The Area Under the Curve (AUC) for Deeplasmid reached 0.93. To be consistent with the standard definition of AUC we have forced predictions to be binary by reducing the standard deviation σ to 0 and setting threshold θ to 0.5. The predictions made by Deeplasmid are averaged over 12 models (Materials and Methods). Fig. 2b shows prediction accuracy individually for each of the 12 models over the ‘validation’ data segment. There is a high agreement between models.

### Feature significance

We also evaluated the significance of various sets of features by calculating the decrease in average AUC of the 12 models on the training data set after setting the features to zero values (thus knocking them out). We retrained the model on various combinations of features. Most of the runs where features were removed resulted in a mean AUC drop more than 3 standard deviations away from the mean AUC achieved when using all features. The use of all features resulted in a mean AUC of 0.897 with a relatively small standard deviation (over 12 runs) of 0.0026. The mean AUC dropped by 5% to 0.847 after removing the hits to plasmid and chromosome-specific genes demonstrating the significance of this feature in classification of annotated contigs. When using only hits to plasmid and chromosome-specific genes (in addition to the sequence data) the mean AUC also dropped to 0.8548. Other features, each one separately, such as sequence length, homopolymer-related features (the longest homopolymer and the total number of homopolymers of length>5), and gene density in the scaffold had relatively little contribution. However, we trained the model with all features since removing features translated to an increase in the error of prediction. We provide an analysis of the mean AUCs observed over 12 models with various feature sets (Table S3). We conclude that using curated biological information provides a clear advantage over previous algorithms, which only used “physical” features of the sequences (such as gc content and scaffold length).

### Testing Deeplasmid model on independent dataset: isolate genomes from IMG database

To test Deeplasmid, we used an independent dataset retrieved from IMG database (48, 49). We downloaded the sequences of 1,834 isolate genomes that have at least one replicon annotated as plasmid. This set included a total of 6820 scaffolds and contigs, with 3093 of them annotated as plasmids and 3727 annotated as chromosomes. Figure 2C shows the ROC curve for the IMG test dataset, achieving an AUC of 0.9344. This suggests that the trained Deeplasmid model is applicable widely, and does not suffer from overfitting. Figure 3 shows the counts of plasmid and chromosomal scaffolds assigned a certain score by Deeplasmid. Setting the threshold for separating the two classes at 0.5, the precision or purity of the predicted positive class (plasmid sequences) is 94%. On the other hand, recall is 77% (details in Suppl. Info) indicating that the DL model missed some plasmids, classifying them as chromosomal fragments.

Similar to our training methodology, the prediction was done on scaffolds of length 1K-330K bases, while scaffolds outside this length range were classified as either too long or too short. There are also 63 ambiguous predictions, which are split between the chromosomal and plasmid classes.

Next, we used this dataset to compare the performance of our DL model to the comparable state-of-the-art programs PlasFlow [16] and cBar [15] (Table 2). Deeplasmid achieved a precision of 0.94 on this dataset (compared to 0.68 for both of the other tools), with a recall comparable to PlasFlow and cBar. Overall, the F1-score, defined as 2·P·R/(P+R), for Deeplasmid was 0.85, which is higher than either PlasFlow (0.69) or cBar (0.73). This comparison demonstrates that addition of biological features vastly improves the performance of programs with otherwise similar inputs and goals.

**Table 2.** Comparison of Deeplasmid (“*D”*) to PLASFlow (“*P”*) to cBar (“*C”*) on the IMG database. With Deeplasmid the purity of the predicted plasmid sequences is 94% (*precision*) and 77% of known plasmids are predicted correctly (*recall*).

|  | Classified as Plasmid |  |  | Classified as Chromosome |  |  | Classified Ambiguous |  |  | Prec (TP/TP+FP) |  |  | Rec (TP/TP+FN) |  |  |
| --- | --- | --- | --- | --- | --- | --- | --- | --- | --- | --- | --- | --- | --- | --- | --- |
|  | *D* | *P* | *C* | *D* | *P* | *C* | *D* | *P* | *C* | *D* | *P* | *C* | *D* | *P* | *C* |
| **Plasmid** | 2264 | 2213 | 2519 | 599 | 229 | 581 | 48 | 658 | 0 | 0.94 | 0.68 | 0.68 | 0.77 | 0.71 | 0.81 |
| **Chromo.** | 140 | 1046 | 1190 | 1687 | 1722 | 2537 | 15 | 959 | 0 | **F1 score:** |  |  | 0.85 | 0.69 | 0.73 |

The runtimes were taken on a Cray XC40 supercomputer (5 Intel Xeon “Haswell” nodes with 120GB, 16 cores). The training runtime was 12.94 hours for the ACLAME and RefSeq.microbial dataset with 41K sequences. For 30 epochs that translates to 26 minutes per epoch. The prediction runtime was <2 seconds per scaffold or under two minutes for a microbial genome assembly, assuming a typical microbial genome assembly contains 1-60 scaffolds. This satisfies the scalability requirement for an automated plasmid finding tool, highlighting Deeplasmid’s potential for widespread application on large-scale genomic and metagenomic data.

### Experimental validation of a new plasmid based on Deeplasmid prediction

In order to demonstrate Deeplasmid’s ability to predict plasmids in biological samples, we focused on the fish pathogen *Yersinia ruckeri* ATCC 29473 (IMG genome ID: 2609460118). Until this work, this strain had only been sequenced by 454 and Illumina short read sequencers, resulting in 15 linear scaffolds and presumably lacking a plasmid (50). Using comparative genomics, researchers studying a similar bacterial strain inferred that *Yersinia ruckeri* ATCC 29473 may encode a plasmid, but no long read sequencing was performed to confirm this (51). The 15 linear scaffolds of *Yersinia ruckeri* ATCC 29473 were then used as input to Deeplasmid, and labeled as either plasmid or chromosomal. One long scaffold of 57Kbp (IMG scaffold 112, Figure 4A) got a Deeplasmid score of 0.8 strongly suggesting that it is derived from a plasmid. In contrast, other scaffolds were either very long (e.g. IMG scaffold 103 of 1.6 Mbp) or received Deeplasmid score below 0.3, suggesting that they derive from a bacterial chromosome. To validate these labels, we grew *Y. ruckeri* bacteria in the lab, extracted DNA, and sequenced it with a long read sequencing method (Oxford Nanopore Technology). In contrast to the previous short read methods, we were able to find large circular pieces of DNA. We found a ∼3.7 Mbp chromosome, and a ∼102 kbp plasmid. Beyond the fact that it is circular, we are confident the latter piece of DNA is a plasmid since it has very high similarity to a known *Yersinia* plasmid (pYR3; Genbank: LN681230.1).

Upon mapping the linear scaffolds onto the newly sequenced circular DNA fragments, we indeed find that predicted plasmid fragments map to the 102 kbp plasmid, demonstrating the predictive power of Deeplasmid and its ability to detect large plasmids in genomic data (Figure 4A). We note that a number of the linear scaffolds for this genome did not undergo Deeplasmid prediction due to their large size (>330kb). Exactly because of their large size, they are assumed to be chromosomal in origin. However, those within the size range of Deeplasmid functionality were largely predicted to be chromosomal in origin, along with some misclassifications (Figure 4).

Looking at the functions of the genes carried on the plasmid, we see many genes previously found on similar plasmids (51), and that have plasmid-related functions. We identified Type IV pilus genes, which may be used for transfer of plasmid from one cell to another (52), or possibly used as a virulence factor (51). Also encoded on the plasmid is the Type IV Secretion System, which may also be involved in plasmid transfer (53) and/or virulence (51). We also detected on the plasmid is RelE, a toxin which is commonly found in plasmid addiction systems (54). Furthermore, we found mobilization genes like transposon genes, integrases, and DNA recombinases. Overall, we conclude that this is a *bona fide* plasmid based on its circularity, separation from the main chromosome, similarity to known plasmids, and plasmidic gene content.

## Discussion and Conclusion

In this work we provide an accurate algorithm to classify assembled contigs or scaffolds generated by any sequencing platform and assembly algorithm, as parts of plasmids and chromosomes using a deep learning approach. By training deep learning models on the specific features of plasmids and chromosomes, we have shown that it is possible to efficiently separate plasmids from chromosomal sequences. While physical sequence features can be used to predict if a sequence is of plasmid or chromosome origin, the DNA sequence itself improves the deep learning models by keeping a memory of what came earlier in the sequence. The reason is that statistical features of sequence composition, such as GC content or oligonucleotide profiles, fail to capture the nucleotide composition over the length of the sequence, as explained in earlier work (8). One of the reasons why our Deeplasmid model has better precision at predicting plasmids than other methods is that it averages predictions over multiple 300bp windows sampled over the length of the sequence, instead of analyzing contigs and scaffolds as a single DNA molecule. As a result, each prediction for a 300bp sequence contributes to the overall result. Additionally, we complemented sequences with extracted feature data that enhance the prediction accuracy. The plasmid-specific ORIs, chromosome-specific genes and GC content serve as essential data features. Averaging the predictions over many 300-base sequences and including biologically meaningful features resulted in a better ability to classify a sequence as plasmidic or chromosomal.

Just like other methods that rely on the nucleotide composition signatures of plasmids and chromosomes, whether as hidden or extracted features, Deeplasmid is likely to have problems with very short sequences for which it may have trouble obtaining a proper sequence signature. For this reason we limited the sequences used in the model construction to those of minimum 1Kb length. There are also very few existing plasmids that are larger than 330 Kbp and therefore we could not train our algorithm on these megaplasmids.

Prediction of plasmids is complicated in large genomic datasets with possible chromosomal integrations of plasmids. Current challenges in the field of plasmid identification include discriminating chromosomes that resemble plasmid sequences or plasmids with chromosomal replication genes. Another challenge is to achieve high prediction accuracy on unknown or understudied microbial lineages that may contain exotic plasmids, since any machine learning tool will be trained with the current knowledge.

Deeplasmid predicts plasmids with a low false positive error rate using only an assembled fasta file as input. It can identify both circular and linear plasmids. The output is a per-scaffold classification of chromosomal, plasmid, or ambiguous contig, along with a score representing the confidence of the prediction. Although the default probability threshold for separation of classes was set at 0.5 based on our benchmarking, users can specify their own filtering cutoffs. Deeplasmid out-performed other available tools in terms of accuracy for single microbial assembly plasmidome analyses. Moreover, we provided experimental evidence for a new plasmid that was predicted using Deeplasmid. A future research direction is to employ Deeplasmid for large-scale identification of plasmids in large-scale metagenomic data from different environments or to uncover novel plasmid-borne antimicrobial resistance genes or novel microbial genes that are horizontally transferred via plasmids. This and myriad other high-impact applications of Deeplasmid are possible due to its fast running time and scalability.

## Supplementary Information

The Supplementary Information file includes Deeplasmid running instructions, comments on features used in chromosome/plasmid classification, Code Repository, Software Design, Deep learning model architecture, AUC per fold on the training dataset, ROC curve for the IMG test dataset, and others.

## Acknowledgements

We thank NERSC for providing us with high-performance computing support throughout this study. This research used resources of the National Energy Research Scientific Computing Center (NERSC), a U.S. Department of Energy Office of Science User Facility operated under Contract No. DE-AC02-05CH11231. The work conducted by the U.S. Department of Energy Joint Genome Institute, a DOE Office of Science User Facility, is supported by the Office of Science of the U.S. Department of Energy under Contract No. DE-AC02-05CH11231.

AL is generously supported by the Israeli Science Foundation (Grants #1535/20, #3300/20), Alon Fellowship of the Israeli council of higher education, The Hebrew University - University of Illinois Urbana-Champaign seed grant, the Israeli Ministry of Agriculture (Grant 12-12-0002), and ICA in Israel. AMG is generously supported by the Kaete Klausner Scholarship and was supported by a scholarship from the Israeli Ministry of Aliyah and Integration. WBA was supported by SJSU’s Faculty startup fund.

We thank Dr. Yasuo Yoshikuni for providing us with *Yersinia ruckeri* ATCC 29473 strain.

