## Supplemental Information for "Deeplasmid: Deep learning accurately separates plasmids from bacterial chromosomes"

### Deeplasmid - Supplementary Information

For questions contact: Bill Andreopoulos, or Asaf Levy

#### Table of Contents:

Instructions to run on a Mac or Linux-based computer:
- Example commands for run on Mac with Docker (10/08/2020):

Building the Docker image
- First issue - include Prodigal in the built container image:
- Second issue - bbmap needs to be in the built Docker image.
- Third issue - the plasmid trained models need to be placed in the Docker image since they are not in the code repository

Features used in chromosome/plasmid classification

Code Repository

Software Design

Deep learning model architecture

AUC per fold on the training dataset

ROC curve for the IMG test dataset

AUC per k-fold and average for combinations of features

Difference of features between 2 classes

#### Instructions to run on a Mac or Linux-based computer:

The deeplasmid software runs on Mac with docker. After installing docker, please run:

```
docker login
docker pull billandreo/deeplasmid
docker run -it  -v /path/to/input/fasta:/srv/jgi-
ml/classifier/dl/in.fasta  -v  /path/to/output/directory:/srv/jgi-
ml/classifier/dl/outdir   billandreo/deeplasmid
feature_DL_plasmid_predict.sh  in.fasta outdir
```

Change the blue highlighted paths above to point to the absolute paths for your input fasta and the output directories. An example run:

```
docker run -it  -v /Users/X/Downloads/jgi-
ml/classifier/dl/temp/649989979.fna:/srv/jgi-
```

```
ml/classifier/dl/in.fasta  -v  /Users/X/Downloads/jgi-
ml/classifier/dl/temp/649989979.fna.OUT:/srv/jgi-
ml/classifier/dl/outdir   billandreo/deeplasmid
feature_DL_plasmid_predict.sh  in.fasta outdir
```

##### Example commands for run on Mac with Docker (10/08/2020):

The two parameters that a user needs to set when running with Docker are the path to input fasta and path to output directory:

```
Williams-MBP:dl andreopo$ pwd
/Users/andreopo/Downloads/jgi-ml_DOCKER_CLEANEDUP/classifier/dl

Williams-MBP:dl andreopo$ docker run -it  -v
/Users/andreopo/Downloads/jgi-
ml_COMMITTED/classifier/dl/temp/649989979.fna:/srv/jgi-
ml/classifier/dl/in.fasta  -v  /Users/andreopo/Downloads/jgi-
ml_DOCKER_CLEANEDUP/classifier/dl/temp/649989979.fna.OUT:/srv/jgi-
ml/classifier/dl/outdir   billandreo/deeplasmid
feature_DL_plasmid_predict.sh  in.fasta outdir
```

A user can check the predictions in the predictions.txt output file:

```
Williams-MBP:dl andreopo$ cat
temp/649989979.fna.OUT/outPR.20201008_211704/predictions.txt
name,pred,conf
NZ_ADHJ01000001,LONGER_330000.0,1
NZ_ADHJ01000014,LONGER_330000.0,1
NZ_ADHJ01000017,LONGER_330000.0,1
NZ_ADHJ01000025,LONGER_330000.0,1
NZ_ADHJ01000027,SHORTER_1000.0,1
NZ_ADHJ01000030,SHORTER_1000.0,1
NZ_ADHJ01000032,SHORTER_1000.0,1
NZ_ADHJ01000037,LONGER_330000.0,1
NZ_ADHJ01000051,SHORTER_1000.0,1
nz_adhj01000006 paenibacillus vortex v453 cnt_pvor1000006, whole
genome shotgun sequence.,GENOME,0.019 +/- 0.001
nz_adhj01000002 paenibacillus vortex v453 cnt_pvor1000002, whole
genome shotgun sequence.,GENOME,0.171 +/- 0.004
nz_adhj01000034 paenibacillus vortex v453 cnt_pvor1000034, whole
genome shotgun sequence.,GENOME,0.134 +/- 0.005
nz_adhj01000013 paenibacillus vortex v453 cnt_pvor1000013, whole
genome shotgun sequence.,GENOME,0.011 +/- 0.001
```

```
nz_adhj01000041 paenibacillus vortex v453 cnt_pvor1000041, whole
genome shotgun sequence.,PLASMID,0.921 +/- 0.001
nz_adhj01000007 paenibacillus vortex v453 cnt_pvor1000007, whole
genome shotgun sequence.,GENOME,0.005 +/- 0.001
...
...
```

#### Building the Docker image

To build the Docker image, please use:

```
docker build -t billandreo/deeplasmid -f Dockerfile.v2  .
```

There are a few issues that need to be addressed when building the Docker image:

##### First issue - include Prodigal in the built container image:

```
git clone https://github.com/hyattpd/Prodigal.git
```

With this command you can enter into the Docker image:

```
docker run -it  billandreo/deeplasmid     /bin/bash
```

Test if prodigal runs with:

```
cd Prodigal/
./Prodigal/prodigal
```

##### Second issue - bbmap needs to be in the built Docker image.

Please check bbmap out from sourceforge into the local dl directory. Commands to place bbmap in the docker image - first place bbmap tar.gz file under the local dl dir:

```
 mv ../../../BBMap_38.73.tar.gz .
 gunzip BBMap_38.73.tar.gz
 tar xvf BBMap_38.73.tar
```

When you build the Docker image with Dockerfile.v2, as shown above, BBMap files will then be placed in the deeplasmid image with the CP command that’s in the Dockerfile.v2

With this command you can enter into the Docker image:

```
docker run -it  billandreo/deeplasmid     /bin/bash
```

This is bbmap version in the Docker container:

```
-rw-r--r--  1 root root 34440192 Jan  1  2020 BBMap_38.73.tar
drwxr-xr-x  9 root root    12288 Jan  4  2020 bbmap
```

##### Third issue - the plasmid trained models need to be placed in the Docker image since they are not in the code repository

For space reasons the models are not in the code repo. Under the local dl directory:

```
cd Plasmid_Models/
scp -r /data/Plasmid_Models/plasmid4z-newfeat5* .
```

Those will then be placed in the deeplasmid image with the CP command that’s in the Dockerfile.v2
Note: for the model you only need .h5 files under Plasmid_Models, which are of a reasonable size. You don’t need the data subdirectories, which are in the GB size range and can cause the Docker container size to become very large. The Plasmid_Models.tar.gz (5MB) can also be sent as email attachment.

#### Features used in chromosome/plasmid classification

**Table S1. A detailed list of the plasmid-specific and chromosome-specific genes and origin-of-replication (ORI) genes is provided in the spreadsheet:**

| **Features** | **Comment** | **Reference** |
|---|---|---|
| Total length of the sequence | Chromosomes tend to be larger than plasmids. | |
| %GC of the sequence | | Rocha and Danchin Trends in Genetics 2002. Base composition bias might result from competition for metabolic resources |
| Longest A homopolymer | | |
| Longest C homopolymer | | |
| Longest G homopolymer | | |
| Longest T homopolymer | | |
| Count of A homopolymer longer than 5 bp | | |
| Count of C homopolymer longer than 5 bp | | |
| Count of G homopolymer longer than 5 bp | | |
| Count of T homopolymer longer than 5 bp | | |
| Contig coding frequency (coding bases/size) | Based on manual inspection of several plasmids, they seem to have lower coding frequency than chromosomes. | |

| | | |
|---|---|---|
| Count of oriV | Plasmid origin of replication | Mei, J., S. Benashski, and W. Firshein, Interactions of the origin of replication (oriV) and initiation proteins (TrfA) of plasmid RK2 with submembrane domains of Escherichia coli. J Bacteriol, 1995. 177(23): p. 6766-72. |
| Count of origin of replication of different plasmids from gram+ and gram- bacteria | Matches to genbank AB015179.1, DQ459477.1, AF239690.1, AF239689.1, X12587.1, EF546766.1, DQ439974.1, V00270.1, Y08803.1, Y08791.1, Y08790.1, X70131.1, V01374.1, V00327.1 | |
| Count of TrfA gene | Plasmid gene | Fang, F.C. and D.R. Helinski, Broad-host-range properties of plasmid RK2: importance of overlapping genes encoding the plasmid replication initiation protein TrfA. J Bacteriol, 1991. 173(18): p. 5861-8. |
| Count of DNA replication initiator gene (DnaA) | Chromosomal gene | Kimelman, A., A. Levy, H. Sberro, S. Kidron, A. Leavitt, G. Amitai, . . . R. Sorek, A vast collection of microbial genes that are toxic to bacteria. Genome Res, 2012. 22(4): p. 802-9. |
| Count of oriC | Chromosomal origin of replication | |
| Count of Rep gene | Plasmid gene | Post□transcriptional control of expression of the repA gene of plasmid R1 mediated by a small RNA molecule. |
| Count of COG0018 | Chromosomal essential gene | Grazziotin, A.L., N.M. Vidal, and T.M. Venancio, Uncovering major genomic features of essential genes in Bacteria and a methanogenic Archaea. FEBS J, 2015. 282(17): p. 3395-3411, 40, and Tazzyman, S.J. and S. Bonhoeffer, Why There Are No Essential Genes on Plasmids. Mol Biol Evol, 2015. 32(12): p. 3079-88. |

| | | |
|---|---|---|
| Count of COG0008 | Chromosomal essential gene | Grazziotin, A.L., N.M. Vidal, and T.M. Venancio, Uncovering major genomic features of essential genes in Bacteria and a methanogenic Archaea. FEBS J, 2015. 282(17): p. 3395-3411, 40, and Tazzyman, S.J. and S. Bonhoeffer, Why There Are No Essential Genes on Plasmids. Mol Biol Evol, 2015. 32(12): p. 3079-88, Sorek, R., Y. Zhu, C.J. Creevey, M.P. Francino, P. Bork, and E.M. Rubin, Genome-wide experimental determination of barriers to horizontal gene transfer. Science, 2007. 318(5855): p. 1449-52. |
| Count of COG0124 | Chromosomal essential gene | Grazziotin, A.L., N.M. Vidal, and T.M. Venancio, Uncovering major genomic features of essential genes in Bacteria and a methanogenic Archaea. FEBS J, 2015. 282(17): p. 3395-3411, 40, and Tazzyman, S.J. and S. Bonhoeffer, Why There Are No Essential Genes on Plasmids. Mol Biol Evol, 2015. 32(12): p. 3079-88. |
| Count of COG0495 | Chromosomal essential gene | Grazziotin, A.L., N.M. Vidal, and T.M. Venancio, Uncovering major genomic features of essential genes in Bacteria and a methanogenic Archaea. FEBS J, 2015. 282(17): p. 3395-3411, 40, and Tazzyman, S.J. and S. Bonhoeffer, Why There Are No Essential Genes on Plasmids. Mol Biol Evol, 2015. 32(12): p. 3079-88. |
| Count of COG0442 | Chromosomal essential gene | Grazziotin, A.L., N.M. Vidal, and T.M. Venancio, Uncovering major genomic features of essential genes in Bacteria and a methanogenic Archaea. FEBS J, 2015. 282(17): p. 3395-3411, 40, and Tazzyman, S.J. and S. Bonhoeffer, Why There Are No |

| | | |
|---|---|---|
| | | Essential Genes on Plasmids. Mol Biol Evol, 2015. 32(12): p. 3079-88. |
| Count of COG0172 | Chromosomal essential gene | Grazziotin, A.L., N.M. Vidal, and T.M. Venancio, Uncovering major genomic features of essential genes in Bacteria and a methanogenic Archaea. FEBS J, 2015. 282(17): p. 3395-3411, 40, and Tazzyman, S.J. and S. Bonhoeffer, Why There Are No Essential Genes on Plasmids. Mol Biol Evol, 2015. 32(12): p. 3079-88. |
| Count of COG0090 | Chromosomal essential and uncloanble gene | Grazziotin, A.L., N.M. Vidal, and T.M. Venancio, Uncovering major genomic features of essential genes in Bacteria and a methanogenic Archaea. FEBS J, 2015. 282(17): p. 3395-3411, 40, and Tazzyman, S.J. and S. Bonhoeffer, Why There Are No Essential Genes on Plasmids. Mol Biol Evol, 2015. 32(12): p. 3079-88. Sorek, R., Y. Zhu, C.J. Creevey, M.P. Francino, P. Bork, and E.M. Rubin, Genome-wide experimental determination of barriers to horizontal gene transfer. Science, 2007. 318(5855): p. 1449-52. |
| Count of COG0087 | Chromosomal essential and unclonable gene | Grazziotin, A.L., N.M. Vidal, and T.M. Venancio, Uncovering major genomic features of essential genes in Bacteria and a methanogenic Archaea. FEBS J, 2015. 282(17): p. 3395-3411, 40, and Tazzyman, S.J. and S. Bonhoeffer, Why There Are No Essential Genes on Plasmids. Mol Biol Evol, 2015. 32(12): p. 3079-88, Sorek, R., Y. Zhu, C.J. Creevey, M.P. Francino, P. Bork, and E.M. Rubin, Genome-wide experimental determination of barriers to horizontal gene transfer. Science, 2007. 318(5855): p. 1449- |

| | | |
|---|---|---|
| | | 52. |
| Count of COG0088 | Chromosomal essential and unclonable gene | Grazziotin, A.L., N.M. Vidal, and T.M. Venancio, Uncovering major genomic features of essential genes in Bacteria and a methanogenic Archaea. FEBS J, 2015. 282(17): p. 3395-3411, 40, and Tazzyman, S.J. and S. Bonhoeffer, Why There Are No Essential Genes on Plasmids. Mol Biol Evol, 2015. 32(12): p. 3079-88, Sorek, R., Y. Zhu, C.J. Creevey, M.P. Francino, P. Bork, and E.M. Rubin, Genome-wide experimental determination of barriers to horizontal gene transfer. Science, 2007. 318(5855): p. 1449-52. |
| Count of COG0097 | Chromosomal essential gene | Grazziotin, A.L., N.M. Vidal, and T.M. Venancio, Uncovering major genomic features of essential genes in Bacteria and a methanogenic Archaea. FEBS J, 2015. 282(17): p. 3395-3411, 40, and Tazzyman, S.J. and S. Bonhoeffer, Why There Are No Essential Genes on Plasmids. Mol Biol Evol, 2015. 32(12): p. 3079-88. |
| Count of COG0102 | Chromosomal essential gene | Grazziotin, A.L., N.M. Vidal, and T.M. Venancio, Uncovering major genomic features of essential genes in Bacteria and a methanogenic Archaea. FEBS J, 2015. 282(17): p. 3395-3411, 40, and Tazzyman, S.J. and S. Bonhoeffer, Why There Are No Essential Genes on Plasmids. Mol Biol Evol, 2015. 32(12): p. 3079-88. |
| Count of COG0092 | Chromosomal essential gene | Grazziotin, A.L., N.M. Vidal, and T.M. Venancio, Uncovering major genomic features of essential genes in Bacteria and a methanogenic Archaea. FEBS J, 2015. 282(17): p. 3395-3411, 40, |

| | | |
|---|---|---|
| | | and Tazzyman, S.J. and S. Bonhoeffer, Why There Are No Essential Genes on Plasmids. Mol Biol Evol, 2015. 32(12): p. 3079-88. |
| Count of COG0522 | Chromosomal essential gene | Grazziotin, A.L., N.M. Vidal, and T.M. Venancio, Uncovering major genomic features of essential genes in Bacteria and a methanogenic Archaea. FEBS J, 2015. 282(17): p. 3395-3411, 40, and Tazzyman, S.J. and S. Bonhoeffer, Why There Are No Essential Genes on Plasmids. Mol Biol Evol, 2015. 32(12): p. 3079-88. |
| Count of COG0098 | Chromosomal essential gene | Grazziotin, A.L., N.M. Vidal, and T.M. Venancio, Uncovering major genomic features of essential genes in Bacteria and a methanogenic Archaea. FEBS J, 2015. 282(17): p. 3395-3411, 40, and Tazzyman, S.J. and S. Bonhoeffer, Why There Are No Essential Genes on Plasmids. Mol Biol Evol, 2015. 32(12): p. 3079-88. |
| Count of COG0202 | Chromosomal essential gene | Grazziotin, A.L., N.M. Vidal, and T.M. Venancio, Uncovering major genomic features of essential genes in Bacteria and a methanogenic Archaea. FEBS J, 2015. 282(17): p. 3395-3411, 40, and Tazzyman, S.J. and S. Bonhoeffer, Why There Are No Essential Genes on Plasmids. Mol Biol Evol, 2015. 32(12): p. 3079-88. |
| Count of COG0592 | Chromosomal essential and unclonable gene | Grazziotin, A.L., N.M. Vidal, and T.M. Venancio, Uncovering major genomic features of essential genes in Bacteria and a methanogenic Archaea. FEBS J, 2015. 282(17): p. 3395-3411, 40, and Tazzyman, S.J. and S. Bonhoeffer, Why There Are No Essential Genes on Plasmids. Mol |

| | | |
|---|---|---|
| | | Biol Evol, 2015. 32(12): p. 3079-88. |
| Count of COG0037 | Chromosomal essential gene | Grazziotin, A.L., N.M. Vidal, and T.M. Venancio, Uncovering major genomic features of essential genes in Bacteria and a methanogenic Archaea. FEBS J, 2015. 282(17): p. 3395-3411, 40, and Tazzyman, S.J. and S. Bonhoeffer, Why There Are No Essential Genes on Plasmids. Mol Biol Evol, 2015. 32(12): p. 3079-88. |
| Count of COG0201 | Chromosomal essential gene | Grazziotin, A.L., N.M. Vidal, and T.M. Venancio, Uncovering major genomic features of essential genes in Bacteria and a methanogenic Archaea. FEBS J, 2015. 282(17): p. 3395-3411, 40, and Tazzyman, S.J. and S. Bonhoeffer, Why There Are No Essential Genes on Plasmids. Mol Biol Evol, 2015. 32(12): p. 3079-88. |
| Count of COG0552 | Chromosomal essential gene | Grazziotin, A.L., N.M. Vidal, and T.M. Venancio, Uncovering major genomic features of essential genes in Bacteria and a methanogenic Archaea. FEBS J, 2015. 282(17): p. 3395-3411, 40, and Tazzyman, S.J. and S. Bonhoeffer, Why There Are No Essential Genes on Plasmids. Mol Biol Evol, 2015. 32(12): p. 3079-88. |
| Count of COG0462 | Chromosomal essential gene | Grazziotin, A.L., N.M. Vidal, and T.M. Venancio, Uncovering major genomic features of essential genes in Bacteria and a methanogenic Archaea. FEBS J, 2015. 282(17): p. 3395-3411, 40, and Tazzyman, S.J. and S. Bonhoeffer, Why There Are No Essential Genes on Plasmids. Mol Biol Evol, 2015. 32(12): p. 3079-88. |

| | | |
|---|---|---|
| Count of COG0593 | Chromosomal unclonable gene | Sorek, R., Y. Zhu, C.J. Creevey, M.P. Francino, P. Bork, and E.M. Rubin, Genome-wide experimental determination of barriers to horizontal gene transfer. Science, 2007. 318(5855): p. 1449-52. |
| Count of COG0776 | Chromosomal unclonable gene | Sorek, R., Y. Zhu, C.J. Creevey, M.P. Francino, P. Bork, and E.M. Rubin, Genome-wide experimental determination of barriers to horizontal gene transfer. Science, 2007. 318(5855): p. 1449-52. |
| Count of COG0592 | Chromosomal unclonable gene | Sorek, R., Y. Zhu, C.J. Creevey, M.P. Francino, P. Bork, and E.M. Rubin, Genome-wide experimental determination of barriers to horizontal gene transfer. Science, 2007. 318(5855): p. 1449-52. |
| Count of COG0206 | Chromosomal unclonable gene | Sorek, R., Y. Zhu, C.J. Creevey, M.P. Francino, P. Bork, and E.M. Rubin, Genome-wide experimental determination of barriers to horizontal gene transfer. Science, 2007. 318(5855): p. 1449-52. |
| Count of COG0234 | Chromosomal unclonable gene | Sorek, R., Y. Zhu, C.J. Creevey, M.P. Francino, P. Bork, and E.M. Rubin, Genome-wide experimental determination of barriers to horizontal gene transfer. Science, 2007. 318(5855): p. 1449-52. |
| Count of COG2885 | Chromosomal unclonable gene | Sorek, R., Y. Zhu, C.J. Creevey, M.P. Francino, P. Bork, and E.M. Rubin, Genome-wide experimental determination of barriers to horizontal gene transfer. Science, 2007. 318(5855): p. 1449-52. |
| Count of COG3203 | Chromosomal unclonable gene | Sorek, R., Y. Zhu, C.J. Creevey, M.P. Francino, P. Bork, and E.M. Rubin, Genome-wide experimental determination of barriers to horizontal gene transfer. Science, 2007. 318(5855): p. 1449-52. |
| Count of COG0048 | Chromosomal unclonable gene | Sorek, R., Y. Zhu, C.J. Creevey, M.P. Francino, P. Bork, and E.M. Rubin, Genome-wide experimental determination of barriers to |

| | | |
|---|---|---|
| | | horizontal gene transfer. Science, 2007. 318(5855): p. 1449-52. |
| Count of COG1278 | Chromosomal unclonable gene | Sorek, R., Y. Zhu, C.J. Creevey, M.P. Francino, P. Bork, and E.M. Rubin, Genome-wide experimental determination of barriers to horizontal gene transfer. Science, 2007. 318(5855): p. 1449-52. |
| Count of COG0052 | Chromosomal unclonable gene | Sorek, R., Y. Zhu, C.J. Creevey, M.P. Francino, P. Bork, and E.M. Rubin, Genome-wide experimental determination of barriers to horizontal gene transfer. Science, 2007. 318(5855): p. 1449-52. |
| Count of COG0050 | Chromosomal unclonable gene | Sorek, R., Y. Zhu, C.J. Creevey, M.P. Francino, P. Bork, and E.M. Rubin, Genome-wide experimental determination of barriers to horizontal gene transfer. Science, 2007. 318(5855): p. 1449-52. |
| Count of COG0228 | Chromosomal unclonable gene | Sorek, R., Y. Zhu, C.J. Creevey, M.P. Francino, P. Bork, and E.M. Rubin, Genome-wide experimental determination of barriers to horizontal gene transfer. Science, 2007. 318(5855): p. 1449-52. |
| Count of COG1028 | Chromosomal unclonable gene | Sorek, R., Y. Zhu, C.J. Creevey, M.P. Francino, P. Bork, and E.M. Rubin, Genome-wide experimental determination of barriers to horizontal gene transfer. Science, 2007. 318(5855): p. 1449-52. |
| Count of COG1959 | Chromosomal unclonable gene | Sorek, R., Y. Zhu, C.J. Creevey, M.P. Francino, P. Bork, and E.M. Rubin, Genome-wide experimental determination of barriers to horizontal gene transfer. Science, 2007. 318(5855): p. 1449-52. |
| Count of COG3104 | Chromosomal unclonable gene | Sorek, R., Y. Zhu, C.J. Creevey, M.P. Francino, P. Bork, and E.M. Rubin, Genome-wide experimental determination of barriers to horizontal gene transfer. Science, 2007. 318(5855): p. 1449-52. |
| Count of COG0739 | Chromosomal unclonable | Sorek, R., Y. Zhu, C.J. Creevey, |

| | | |
|---|---|---|
| | gene | M.P. Francino, P. Bork, and E.M. Rubin, Genome-wide experimental determination of barriers to horizontal gene transfer. Science, 2007. 318(5855): p. 1449-52. |
| Count of COG0594 | Chromosomal unclonable gene | Sorek, R., Y. Zhu, C.J. Creevey, M.P. Francino, P. Bork, and E.M. Rubin, Genome-wide experimental determination of barriers to horizontal gene transfer. Science, 2007. 318(5855): p. 1449-52. |
| Count of COG0774 | Chromosomal unclonable gene | Sorek, R., Y. Zhu, C.J. Creevey, M.P. Francino, P. Bork, and E.M. Rubin, Genome-wide experimental determination of barriers to horizontal gene transfer. Science, 2007. 318(5855): p. 1449-52. |
| Count of COG1077 | Chromosomal unclonable gene | Sorek, R., Y. Zhu, C.J. Creevey, M.P. Francino, P. Bork, and E.M. Rubin, Genome-wide experimental determination of barriers to horizontal gene transfer. Science, 2007. 318(5855): p. 1449-52. |
| Count of COG0716 | Chromosomal unclonable gene | Sorek, R., Y. Zhu, C.J. Creevey, M.P. Francino, P. Bork, and E.M. Rubin, Genome-wide experimental determination of barriers to horizontal gene transfer. Science, 2007. 318(5855): p. 1449-52. |
| Count of COG0799 | Chromosomal unclonable gene | Sorek, R., Y. Zhu, C.J. Creevey, M.P. Francino, P. Bork, and E.M. Rubin, Genome-wide experimental determination of barriers to horizontal gene transfer. Science, 2007. 318(5855): p. 1449-52. |
| Count of COG1734 | Chromosomal unclonable gene | Sorek, R., Y. Zhu, C.J. Creevey, M.P. Francino, P. Bork, and E.M. Rubin, Genome-wide experimental determination of barriers to horizontal gene transfer. Science, 2007. 318(5855): p. 1449-52. |
| Count of COG0089 | Chromosomal unclonable gene | Sorek, R., Y. Zhu, C.J. Creevey, M.P. Francino, P. Bork, and E.M. Rubin, Genome-wide experimental determination of barriers to horizontal gene transfer. Science, |

| | | |
|---|---|---|
| | | 2007. 318(5855): p. 1449-52. |
| Count of COG0849 | Chromosomal unclonable gene | Sorek, R., Y. Zhu, C.J. Creevey, M.P. Francino, P. Bork, and E.M. Rubin, Genome-wide experimental determination of barriers to horizontal gene transfer. Science, 2007. 318(5855): p. 1449-52. |
| Count of COG1686 | Chromosomal unclonable gene | Sorek, R., Y. Zhu, C.J. Creevey, M.P. Francino, P. Bork, and E.M. Rubin, Genome-wide experimental determination of barriers to horizontal gene transfer. Science, 2007. 318(5855): p. 1449-52. |
| Count of COG1399 | Chromosomal unclonable gene | Sorek, R., Y. Zhu, C.J. Creevey, M.P. Francino, P. Bork, and E.M. Rubin, Genome-wide experimental determination of barriers to horizontal gene transfer. Science, 2007. 318(5855): p. 1449-52. |
| Count of COG0236 | Chromosomal unclonable gene | Sorek, R., Y. Zhu, C.J. Creevey, M.P. Francino, P. Bork, and E.M. Rubin, Genome-wide experimental determination of barriers to horizontal gene transfer. Science, 2007. 318(5855): p. 1449-52. |
| Count of COG1414 | Chromosomal unclonable gene | Sorek, R., Y. Zhu, C.J. Creevey, M.P. Francino, P. Bork, and E.M. Rubin, Genome-wide experimental determination of barriers to horizontal gene transfer. Science, 2007. 318(5855): p. 1449-52. |
| Count of COG0332 | Chromosomal unclonable gene | Sorek, R., Y. Zhu, C.J. Creevey, M.P. Francino, P. Bork, and E.M. Rubin, Genome-wide experimental determination of barriers to horizontal gene transfer. Science, 2007. 318(5855): p. 1449-52. |
| Count of COG0527 | Chromosomal unclonable gene | Sorek, R., Y. Zhu, C.J. Creevey, M.P. Francino, P. Bork, and E.M. Rubin, Genome-wide experimental determination of barriers to horizontal gene transfer. Science, 2007. 318(5855): p. 1449-52. |
| Count of COG0102 | Chromosomal unclonable gene | Sorek, R., Y. Zhu, C.J. Creevey, M.P. Francino, P. Bork, and E.M. |

| | | |
|---|---|---|
| | | Rubin, Genome-wide experimental determination of barriers to horizontal gene transfer. Science, 2007. 318(5855): p. 1449-52. |
| Count of COG0361 | Chromosomal unclonable gene | Sorek, R., Y. Zhu, C.J. Creevey, M.P. Francino, P. Bork, and E.M. Rubin, Genome-wide experimental determination of barriers to horizontal gene transfer. Science, 2007. 318(5855): p. 1449-52. |
| Count of COG0568 | Chromosomal unclonable gene | Sorek, R., Y. Zhu, C.J. Creevey, M.P. Francino, P. Bork, and E.M. Rubin, Genome-wide experimental determination of barriers to horizontal gene transfer. Science, 2007. 318(5855): p. 1449-52. |
| Count of COG0764 | Chromosomal unclonable gene | Sorek, R., Y. Zhu, C.J. Creevey, M.P. Francino, P. Bork, and E.M. Rubin, Genome-wide experimental determination of barriers to horizontal gene transfer. Science, 2007. 318(5855): p. 1449-52. |
| Count of COG0199 | Chromosomal unclonable gene | Sorek, R., Y. Zhu, C.J. Creevey, M.P. Francino, P. Bork, and E.M. Rubin, Genome-wide experimental determination of barriers to horizontal gene transfer. Science, 2007. 318(5855): p. 1449-52. |
| Count of COG0845 | Chromosomal unclonable gene | Sorek, R., Y. Zhu, C.J. Creevey, M.P. Francino, P. Bork, and E.M. Rubin, Genome-wide experimental determination of barriers to horizontal gene transfer. Science, 2007. 318(5855): p. 1449-52. |
| Count of COG0545 | Chromosomal unclonable gene | Sorek, R., Y. Zhu, C.J. Creevey, M.P. Francino, P. Bork, and E.M. Rubin, Genome-wide experimental determination of barriers to horizontal gene transfer. Science, 2007. 318(5855): p. 1449-52. |
| Count of COG0261 | Chromosomal unclonable gene | Sorek, R., Y. Zhu, C.J. Creevey, M.P. Francino, P. Bork, and E.M. Rubin, Genome-wide experimental determination of barriers to horizontal gene transfer. Science, 2007. 318(5855): p. 1449-52. |

| | | |
|---|---|---|
| Count of COG0834 | Chromosomal unclonable gene | Sorek, R., Y. Zhu, C.J. Creevey, M.P. Francino, P. Bork, and E.M. Rubin, Genome-wide experimental determination of barriers to horizontal gene transfer. Science, 2007. 318(5855): p. 1449-52. |
| Count of COG0189 | Chromosomal unclonable gene | Sorek, R., Y. Zhu, C.J. Creevey, M.P. Francino, P. Bork, and E.M. Rubin, Genome-wide experimental determination of barriers to horizontal gene transfer. Science, 2007. 318(5855): p. 1449-52. |
| Count of COG0099 | Chromosomal unclonable gene | Sorek, R., Y. Zhu, C.J. Creevey, M.P. Francino, P. Bork, and E.M. Rubin, Genome-wide experimental determination of barriers to horizontal gene transfer. Science, 2007. 318(5855): p. 1449-52. |
| Count of COG1314 | Chromosomal unclonable gene | Sorek, R., Y. Zhu, C.J. Creevey, M.P. Francino, P. Bork, and E.M. Rubin, Genome-wide experimental determination of barriers to horizontal gene transfer. Science, 2007. 318(5855): p. 1449-52. |
| Count of COG0049 | Chromosomal unclonable gene | Sorek, R., Y. Zhu, C.J. Creevey, M.P. Francino, P. Bork, and E.M. Rubin, Genome-wide experimental determination of barriers to horizontal gene transfer. Science, 2007. 318(5855): p. 1449-52. |
| Count of COG0336 | Chromosomal unclonable gene | Sorek, R., Y. Zhu, C.J. Creevey, M.P. Francino, P. Bork, and E.M. Rubin, Genome-wide experimental determination of barriers to horizontal gene transfer. Science, 2007. 318(5855): p. 1449-52. |
| Count of COG3248 | Chromosomal unclonable gene | Sorek, R., Y. Zhu, C.J. Creevey, M.P. Francino, P. Bork, and E.M. Rubin, Genome-wide experimental determination of barriers to horizontal gene transfer. Science, 2007. 318(5855): p. 1449-52. |
| Count of COG0185 | Chromosomal unclonable gene | Sorek, R., Y. Zhu, C.J. Creevey, M.P. Francino, P. Bork, and E.M. Rubin, Genome-wide experimental determination of barriers to |

| | | |
|---|---|---|
| | | horizontal gene transfer. Science, 2007. 318(5855): p. 1449-52. |
| Count of COG0051 | Chromosomal unclonable gene | Sorek, R., Y. Zhu, C.J. Creevey, M.P. Francino, P. Bork, and E.M. Rubin, Genome-wide experimental determination of barriers to horizontal gene transfer. Science, 2007. 318(5855): p. 1449-52. |
| Count of COG2814 | Chromosomal unclonable gene | Sorek, R., Y. Zhu, C.J. Creevey, M.P. Francino, P. Bork, and E.M. Rubin, Genome-wide experimental determination of barriers to horizontal gene transfer. Science, 2007. 318(5855): p. 1449-52. |
| Count of COG1475 | Chromosomal unclonable gene | Sorek, R., Y. Zhu, C.J. Creevey, M.P. Francino, P. Bork, and E.M. Rubin, Genome-wide experimental determination of barriers to horizontal gene transfer. Science, 2007. 318(5855): p. 1449-52. |
| Count of COG0227 | Chromosomal unclonable gene | Sorek, R., Y. Zhu, C.J. Creevey, M.P. Francino, P. Bork, and E.M. Rubin, Genome-wide experimental determination of barriers to horizontal gene transfer. Science, 2007. 318(5855): p. 1449-52. |
| Count of COG0057 | Chromosomal unclonable gene | Sorek, R., Y. Zhu, C.J. Creevey, M.P. Francino, P. Bork, and E.M. Rubin, Genome-wide experimental determination of barriers to horizontal gene transfer. Science, 2007. 318(5855): p. 1449-52. |
| Count of COG0501 | Chromosomal unclonable gene | Sorek, R., Y. Zhu, C.J. Creevey, M.P. Francino, P. Bork, and E.M. Rubin, Genome-wide experimental determination of barriers to horizontal gene transfer. Science, 2007. 318(5855): p. 1449-52. |
| Count of COG4465 | Chromosomal unclonable gene | Sorek, R., Y. Zhu, C.J. Creevey, M.P. Francino, P. Bork, and E.M. Rubin, Genome-wide experimental determination of barriers to horizontal gene transfer. Science, 2007. 318(5855): p. 1449-52. |
| Count of COG0534 | Chromosomal unclonable | Sorek, R., Y. Zhu, C.J. Creevey, |

| | | |
|---|---|---|
| | gene | M.P. Francino, P. Bork, and E.M. Rubin, Genome-wide experimental determination of barriers to horizontal gene transfer. Science, 2007. 318(5855): p. 1449-52. |
| Count of COG0081 | Chromosomal unclonable gene | Sorek, R., Y. Zhu, C.J. Creevey, M.P. Francino, P. Bork, and E.M. Rubin, Genome-wide experimental determination of barriers to horizontal gene transfer. Science, 2007. 318(5855): p. 1449-52. |
| Count of COG0335 | Chromosomal unclonable gene | Sorek, R., Y. Zhu, C.J. Creevey, M.P. Francino, P. Bork, and E.M. Rubin, Genome-wide experimental determination of barriers to horizontal gene transfer. Science, 2007. 318(5855): p. 1449-52. |
| Count of COG0583 | Chromosomal unclonable gene | Sorek, R., Y. Zhu, C.J. Creevey, M.P. Francino, P. Bork, and E.M. Rubin, Genome-wide experimental determination of barriers to horizontal gene transfer. Science, 2007. 318(5855): p. 1449-52. |
| Count of Hok | Plasmid genes, toxin-antixoin | Unterholzner, S.J., B. Poppenberger, and W. Rozhon, Toxin-antitoxin systems: Biology, identification, and application. Mob Genet Elements, 2013. 3(5): p. e26219. |
| Count of TisB | Plasmid genes, toxin-antixoin | Unterholzner, S.J., B. Poppenberger, and W. Rozhon, Toxin-antitoxin systems: Biology, identification, and application. Mob Genet Elements, 2013. 3(5): p. e26219. |
| Count of SymE | Plasmid genes, toxin-antixoin | Unterholzner, S.J., B. Poppenberger, and W. Rozhon, Toxin-antitoxin systems: Biology, identification, and application. Mob Genet Elements, 2013. 3(5): p. e26219. |
| Count of CcdB-CcdA | Plasmid genes, toxin-antixoin | Unterholzner, S.J., B. Poppenberger, and W. Rozhon, Toxin-antitoxin systems: Biology, identification, and application. Mob Genet Elements, 2013. 3(5): p. |

| | | |
|---|---|---|
| | | e26219. |
| Count of ParE-ParD | Plasmid genes, toxin-antixoin | Unterholzner, S.J., B. Poppenberger, and W. Rozhon, Toxin-antitoxin systems: Biology, identification, and application. Mob Genet Elements, 2013. 3(5): p. e26219. |
| Count of MazF-MazE | Plasmid genes, toxin-antixoin | Unterholzner, S.J., B. Poppenberger, and W. Rozhon, Toxin-antitoxin systems: Biology, identification, and application. Mob Genet Elements, 2013. 3(5): p. e26219. |
| Count of Kid-Kis | Plasmid genes, toxin-antixoin | Unterholzner, S.J., B. Poppenberger, and W. Rozhon, Toxin-antitoxin systems: Biology, identification, and application. Mob Genet Elements, 2013. 3(5): p. e26219. |
| Count of HicA-HicB | Plasmid genes, toxin-antixoin | Unterholzner, S.J., B. Poppenberger, and W. Rozhon, Toxin-antitoxin systems: Biology, identification, and application. Mob Genet Elements, 2013. 3(5): p. e26219. |
| Count of RelE-RelB | Plasmid genes, toxin-antixoin | Unterholzner, S.J., B. Poppenberger, and W. Rozhon, Toxin-antitoxin systems: Biology, identification, and application. Mob Genet Elements, 2013. 3(5): p. e26219. |
| Count of VapC-VapB | Plasmid genes, toxin-antixoin | Unterholzner, S.J., B. Poppenberger, and W. Rozhon, Toxin-antitoxin systems: Biology, identification, and application. Mob Genet Elements, 2013. 3(5): p. e26219. |
| Count of Doc-PhD | Plasmid genes, toxin-antixoin | Unterholzner, S.J., B. Poppenberger, and W. Rozhon, Toxin-antitoxin systems: Biology, identification, and application. Mob Genet Elements, 2013. 3(5): p. e26219. |
| Count of RatA-RatB | Plasmid genes, toxin-antixoin | Unterholzner, S.J., B. Poppenberger, and W. Rozhon, |

| | | |
|---|---|---|
| | | Toxin-antitoxin systems: Biology, identification, and application. Mob Genet Elements, 2013. 3(5): p. e26219. |
| Count of HipA-HipB | Plasmid genes, toxin-antixoin | Unterholzner, S.J., B. Poppenberger, and W. Rozhon, Toxin-antitoxin systems: Biology, identification, and application. Mob Genet Elements, 2013. 3(5): p. e26219. |
| Count of ToxN-ToxI | Plasmid genes, toxin-antixoin | Unterholzner, S.J., B. Poppenberger, and W. Rozhon, Toxin-antitoxin systems: Biology, identification, and application. Mob Genet Elements, 2013. 3(5): p. e26219. |
| Count of YeeV-Yeeu | Plasmid genes, toxin-antixoin | Unterholzner, S.J., B. Poppenberger, and W. Rozhon, Toxin-antitoxin systems: Biology, identification, and application. Mob Genet Elements, 2013. 3(5): p. e26219. |
| Count of CptA-CptB | Plasmid genes, toxin-antixoin | Unterholzner, S.J., B. Poppenberger, and W. Rozhon, Toxin-antitoxin systems: Biology, identification, and application. Mob Genet Elements, 2013. 3(5): p. e26219. |
| Count of GhoT-GhoS | Plasmid genes, toxin-antixoin | Unterholzner, S.J., B. Poppenberger, and W. Rozhon, Toxin-antitoxin systems: Biology, identification, and application. Mob Genet Elements, 2013. 3(5): p. e26219. |
| Count of Par gene | Plasmid gene | Gerdes, K., J. Moller-Jensen, and R. Bugge Jensen, Plasmid and chromosome partitioning: surprises from phylogeny. Mol Microbiol, 2000. 37(3): p. 455-66. |
| Count of Psi gene | Plasmid gene | Petrova, V., S. Chitteni-Pattu, J.C. Drees, R.B. Inman, and M.M. Cox, An SOS inhibitor that binds to free RecA protein: the PsiB protein. Mol Cell, 2009. 36(1): p. 121-30. |

| | | |
|---|---|---|
| Count of Tra gene | Plasmid gene, responsible for plasmid transfer | Zatyka, M. and C.M. Thomas, Control of genes for conjugative transfer of plasmids and other mobile elements. FEMS Microbiol Rev, 1998. 21: p. 291-319. |
| Count of Stb gene | Plasmid gene, responsible for plasmid stability | Guynet, C., A. Cuevas, G. Moncalian, and F. de la Cruz, The stb operon balances the requirements for vegetative stability and conjugative transfer of plasmid R388. PLoS Genet, 2011. 7(5): p. e1002073. |
| Count of Mob gene | Plasmid gene | Wang, P., Y. Zhu, Y. Zhang, C. Zhang, J. Xu, Y. Deng, . . . M. Sun, Mob/oriT, a mobilizable site-specific recombination system for unmarked genetic manipulation in Bacillus thuringiensis and Bacillus cereus. Microb Cell Fact, 2016. 15(1): p. 108. |
| Count of relaxase gene | Plasmid gene | Garcillan-Barcia, M.P., M.V. Francia, and F. de la Cruz, The diversity of conjugative relaxases and its application in plasmid classification. FEMS Microbiol Rev, 2009. 33(3): p. 657-87. |
| Count of Type IV pili coupling protein | Plasmid gene | Smillie, C., M.P. Garcillan-Barcia, M.V. Francia, E.P. Rocha, and F. de la Cruz, Mobility of plasmids. Microbiol Mol Biol Rev, 2010. 74(3): p. 434-52. |
| Count of VirB gene | Plasmid gene | Smillie, C., M.P. Garcillan-Barcia, M.V. Francia, E.P. Rocha, and F. de la Cruz, Mobility of plasmids. Microbiol Mol Biol Rev, 2010. 74(3): p. 434-52. |

#### Code Repository

The Sourceforge code repository is at: deeplasmid.sourceforge.io
https://sourceforge.net/projects/deeplasmid/
This repo is mainly intended to store code for the paper and open-source community access.

The classifier/dl directory had the cleaned up code. You may delete or ignore other dirs, like bin and binning dirs.
The **docker** branch contains the docker-specific code that works on both the *Mac (with docker) and Cori (with shifter).* The **master** branch is the version that was deployed natively on Cori (in jgi/lbnl).

#### Software Design

A flowchart of the software design showing the modules and what files get input to the prediction and training:

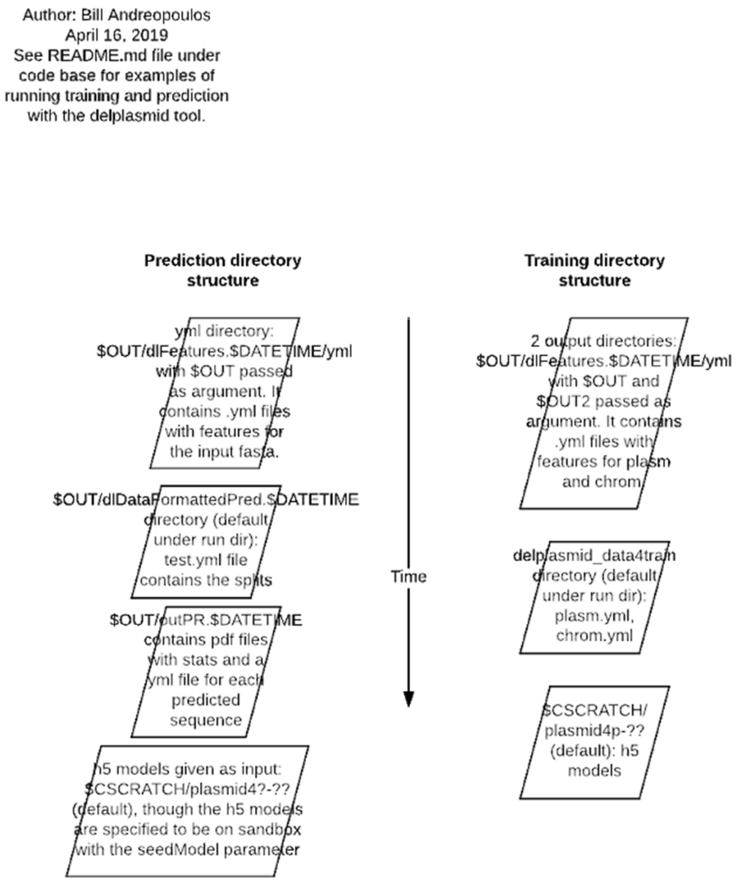


**Figure S1:** The directory structure expected for input to the training and prediction and the output directories produced

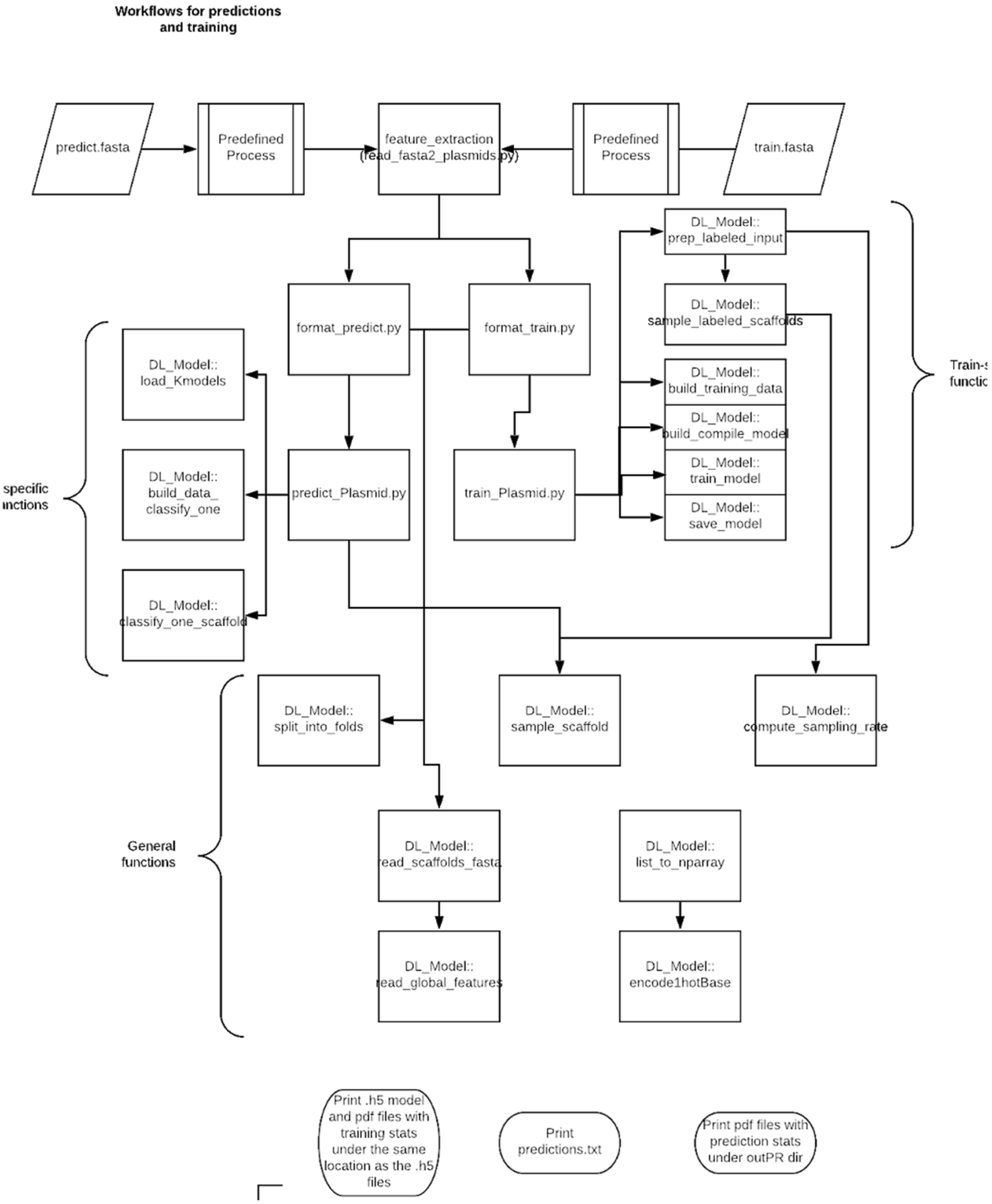


**Figure S2:** The design of the codebase with the main classes, important functions, and call graph

#### Deep learning model architecture

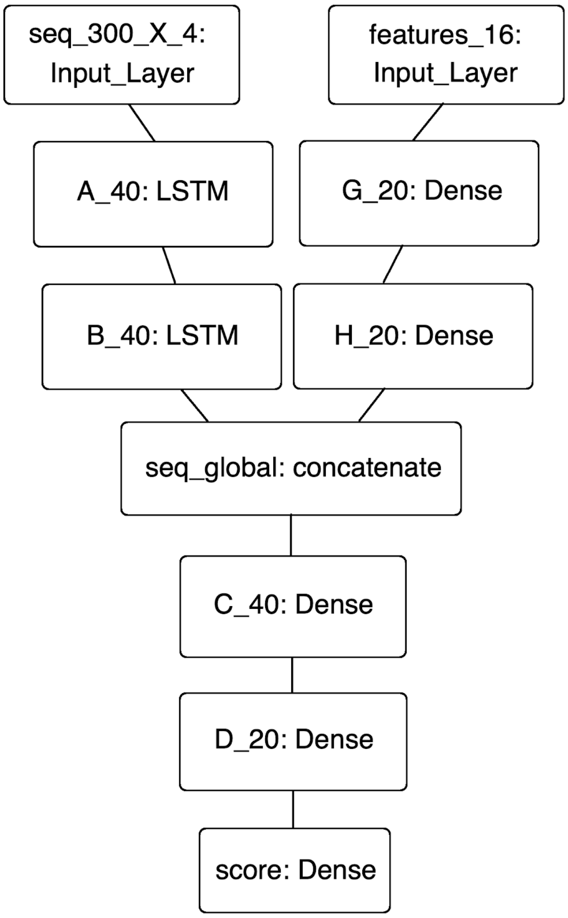


**Figure S3: Model topology of Deeplasmid Neural Network.** Fully connected layers are denoted as 'Dense'. The numbers in the boxes are the sizes of the output features. This model has a total of 22,865 parameters. A dropout rate of 0.1 is applied in-between all layers. Dropout sets weights to zero in order to prevent overfitting and allow the training to generalize to new datasets.

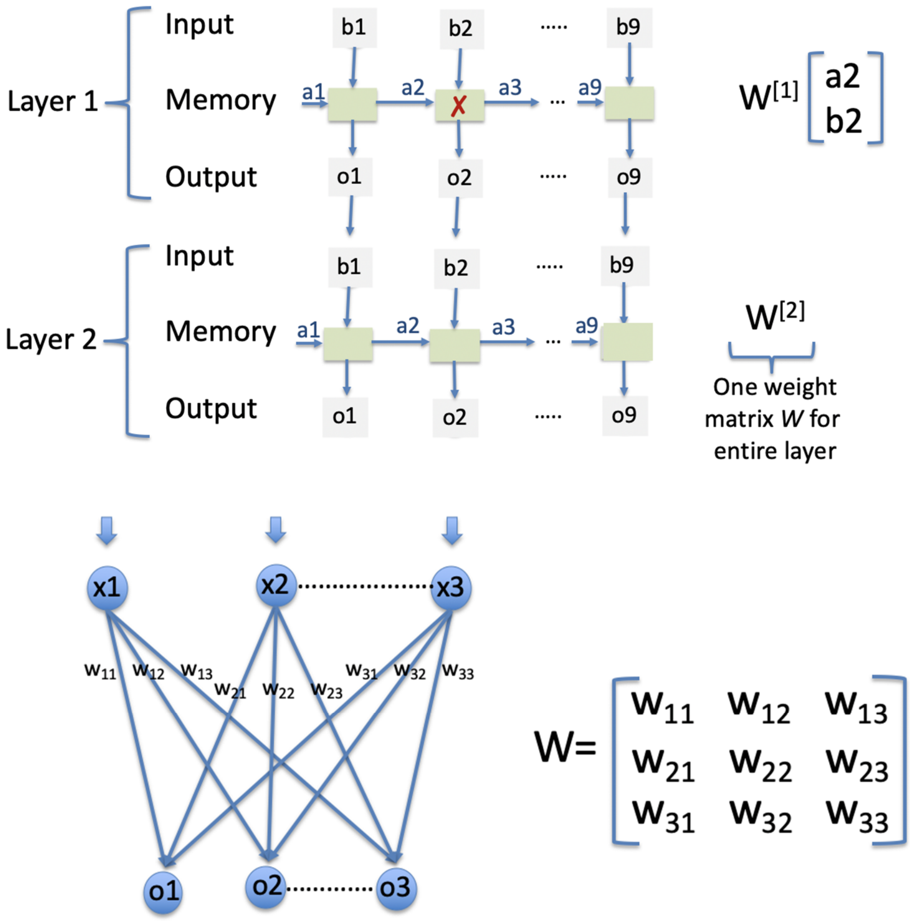


**Figure S4:** A layered LSTM architecture (top), and a basic dense-layer architecture (bottom).

AUC per fold on the training dataset

| fold | AUC |
|---|---|
| 0 | 0.954186 |
| 1 | 0.945295 |
| 2 | 0.964656 |
| 3 | 0.953013 |
| 4 | 0.957194 |
| 5 | 0.963042 |
| 0 | 0.959198 |
| 1 | 0.931794 |
| 2 | 0.968798 |
| 3 | 0.949437 |
| 4 | 0.956179 |
| 5 | 0.953591 |

**Table S2:** We computed the AUC per each of the 12 models (2 models per fold)

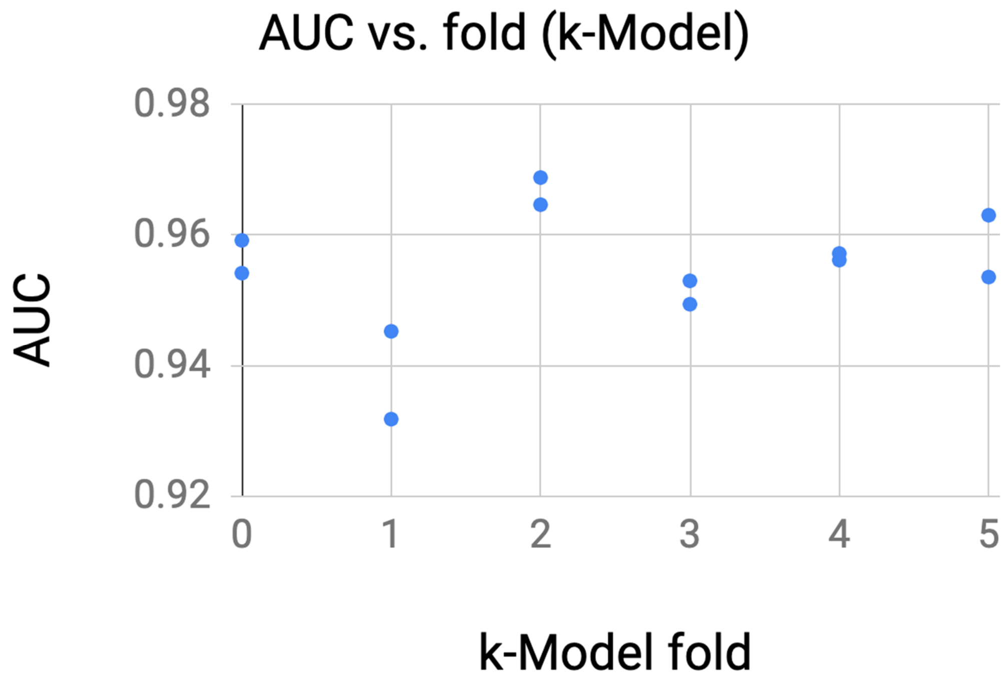

**Figure S5:** The AUC achieved for each k-Fold model

#### ROC curve for the IMG test dataset

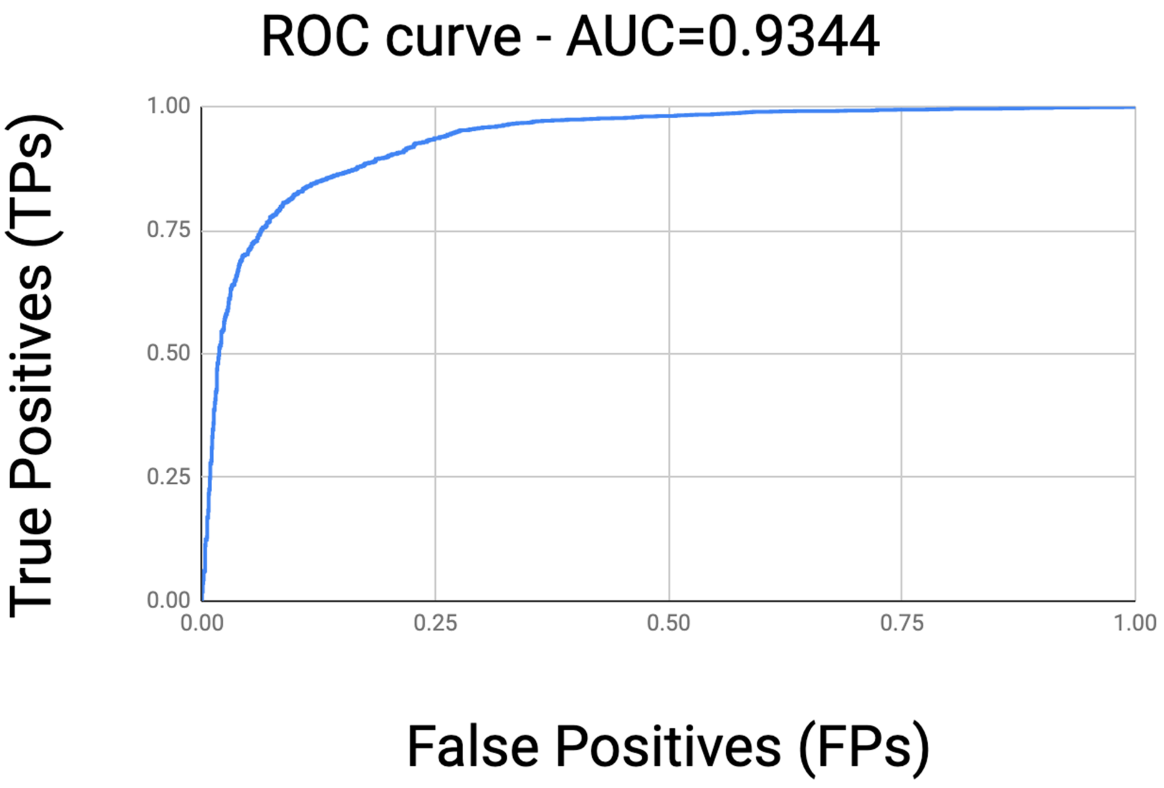


**Figure S6:** The ROC curve (TPs vs. FPs) for the IMG test dataset.

#### AUC per k-fold and average for combinations of features

The spreadsheet with the cross-validation results shown in this table is provided under: https://docs.google.com/spreadsheets/d/1TDPn9uOAnZOBS95dJzUfVtb6mhteTuRFrFcv-_lZ4z4/edit#gid=1266368803

| **AUC** | **1** | **2** | **3** | **4** | **5** | **6** | **7** | **8** | **9** | **10** | **11** | **12** | **AVG** | **STD** |
|---|---|---|---|---|---|---|---|---|---|---|---|---|---|---|
| All features | 0.8889 | 0.8921 | 0.914 | 0.896 | 0.8974 | 0.9035 | 0.8898 | 0.8835 | 0.9124 | 0.8964 | 0.8962 | 0.8944 | **0.89705** | **0.002612** |
| All features_minus_aa_avg_len | 0.8989 | 0.8833 | 0.889 | 0.8818 | 0.8914 | 0.8933 | 0.8889 | 0.8684 | 0.905 | 0.8789 | 0.8888 | 0.9006 | **0.889025** | **0.002917** |
| All features_minus_ | 0.9105 | 0.8707 | 0.9163 | 0.9019 | 0.9038 | 0.8934 | 0.9056 | 0.8843 | 0.9174 | 0.8869 | 0.9064 | 0.8955 | **0.89939** | **0.003993** |

| | | | | | | | | | | | | | |
|---|---|---|---|---|---|---|---|---|---|---|---|---|---|
| len_sequence | | | | | | | | | | | | | |
| All features_minus_gc_content | 0.9068 | 0.8916 | 0.9021 | 0.8779 | 0.8997 | 0.8962 | 0.8952 | 0.8903 | 0.9133 | 0.877 | 0.9028 | 0.8924 | **0.89544** | **0.003093** |
| hits_chrom_plasmid_proteins_ORIs_only | 0.8561 | 0.8529 | 0.8593 | 0.8497 | 0.863 | 0.8544 | 0.8473 | 0.8445 | 0.8637 | 0.8505 | 0.857 | 0.8595 | **0.854825** | **0.001753** |
| All features_minus_hits_chrom_plasmid_proteins_ORIs | 0.8574 | 0.8291 | 0.8411 | 0.8516 | 0.8251 | 0.8557 | 0.8609 | 0.8381 | 0.8542 | 0.8517 | 0.844 | 0.8613 | **0.84751** | **0.003477** |
| All features_minus_gene_count | 0.889 | 0.8789 | 0.9122 | 0.8803 | 0.9057 | 0.8974 | 0.9007 | 0.8786 | 0.9143 | 0.8806 | 0.8952 | 0.891 | **0.89365** | **0.003695** |
| gene_count_aa_avg_len_only | 0.8124 | 0.7841 | | 0.7837 | 0.7841 | 0.8018 | 0.8104 | 0.7679 | 0.821 | 0.7901 | 0.7746 | 0.7771 | **0.79156** | **0.004993** |
| All features_minus_homopolymer | 0.8835 | 0.8735 | 0.9143 | 0.8911 | 0.8967 | 0.8862 | 0.881 | 0.8698 | 0.9088 | 0.8878 | 0.896 | 0.8925 | **0.8901** | **0.003748** |
| gc_content_lensequence_only | 0.7434 | 0.7175 | 0.7439 | 0.7178 | 0.7163 | 0.7497 | 0.7425 | 0.7237 | 0.7324 | 0.7328 | 0.7204 | 0.7406 | **0.73175** | **0.003515** |

**Table S3:** The AUC achieved for various combinations of features shows the best AUC is achieved when keeping all features.

After including the following 5 features the AUC on the IMG test dataset increased by about 7% from 86.91% to 93.97%:

- A binary feature (1/0) if a contig hits any plasmid-specific amino acid sequences (plasmidProt).
- A binary feature (1/0) if a contig hits any plasmid-specific ORI nucleotide sequences (plasmid_originOfReplication.nr.fasta).
- A binary feature (1/0) if a contig hits any chromosome-specific amino acid sequences (these aa sequences are in the .cdhit files, cdhit is a clustering tool).
- number of genes in a contig.
- percent of gene-coding sequence in a contig.

#### Difference of features between genomic DNA (chromosomes) and plasmids

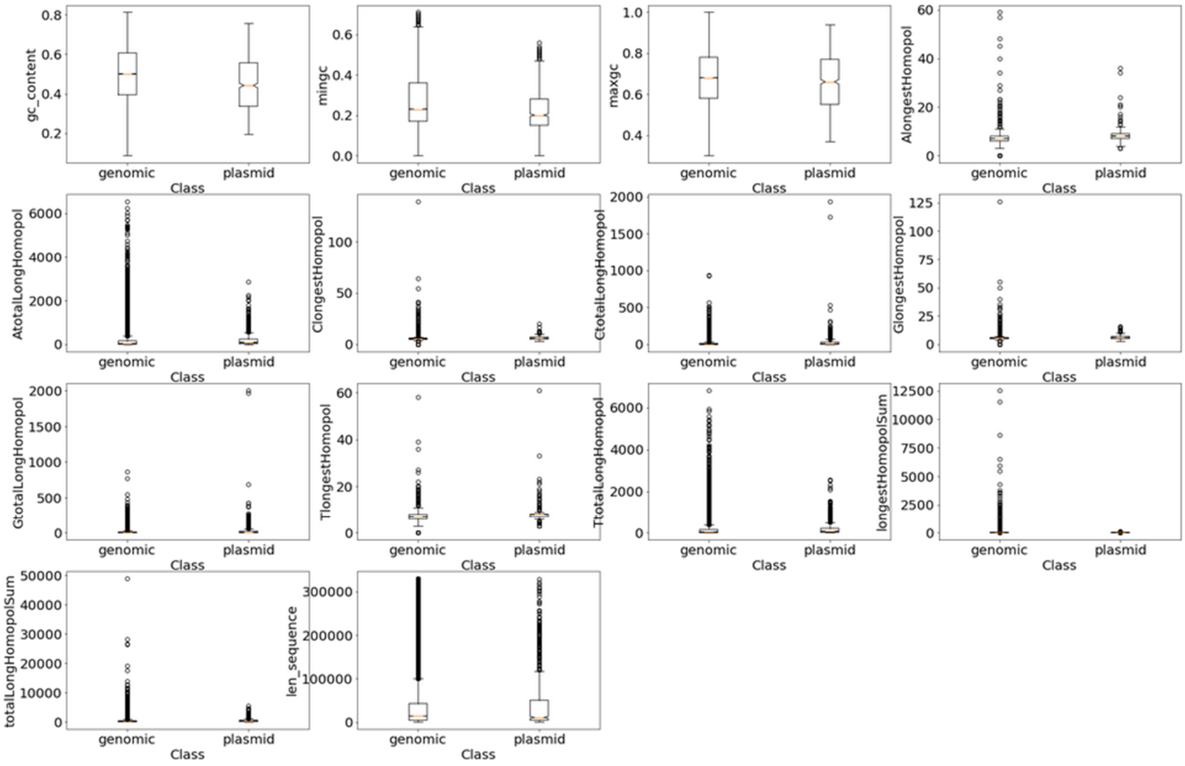


**Figure S7:** The feature values compared between the two classes of chromosomal vs. plasmid
